# Transgenic goats producing an improved version of cetuximab in milk

**DOI:** 10.1101/2020.06.05.137463

**Authors:** Götz Laible, Sally Cole, Brigid Brophy, Paul Maclean, Li How Chen, Dan P. Pollock, Lisa Cavacini, Nathalie Fournier, Christophe De Romeuf, Nicholas C. Masiello, William G. Gavin, David N. Wells, Harry M. Meade

**Affiliations:** AgResearch, Ruakura Research Centre, Hamilton 3240, New Zealand; School of Medical Sciences, University of Auckland, Auckland 1023, New Zealand; LFB-USA, Framingham, MA 01702, United States; LFB Biotechnologies, 59000 Lille, France; MassBiologics of the University of Massachusetts Medical School, Boston, MA 02126 United States

**Author notes:** Corresponding Author: Götz Laible, AgResearch Ruakura Research Centre, Hamilton 3240, New Zealand, Fax +64 7.

**Keywords:** Monoclonal antibody, biosimilar, EGFR, biobetter

## Abstract

Therapeutic monoclonal antibodies (mAbs) represent one of the most important classes of pharmaceutical proteins to treat human diseases. Most are produced in cultured mammalian cells which is expensive, limiting their availability. Goats, striking a good balance between a relatively short generation time and copious milk yield, present an alternative platform for the cost-effective, flexible, large-scale production of therapeutic mAbs. Here, we focused on cetuximab, a mAb against epidermal growth factor receptor, that is commercially produced under the brand name Erbitux and approved for anti-cancer treatments. We generated several transgenic goat lines that produce cetuximab in their milk. Two lines were selected for detailed characterization. Both showed stable genotypes and cetuximab production levels of up to 10g/L. The mAb could be readily purified and showed improved characteristics compared to Erbitux. The goat-produced cetuximab (gCetuximab) lacked a highly immunogenic epitope that is part of Erbitux. Moreover, it showed enhanced binding to CD16 and increased antibody-dependent cell-dependent cytotoxicity compared to Erbitux. This indicates that these goats produce an improved cetuximab version with the potential for enhanced effectiveness and better safety profile compared to treatments with Erbitux. In addition, our study validates transgenic goats as an excellent platform for large-scale production of therapeutic mAbs.

## INTRODUCTION

Monoclonal antibodies (mAbs) represent an important class of biopharmaceutical drugs for the treatment of serious human diseases, with many ranked within the ten top-selling biopharmaceuticals (1). They are commonly produced as recombinant proteins in cultured mammalian cells to ensure proper glycosylation which is often required for full functionality and makes it currently the prevailing and most accepted production system (2, 3). Still, high up-front capital investment for the production facilities and lack of scalability are recognized weaknesses of this technology (4, 5). Dairy animals and production of recombinant therapeutic proteins in milk could provide an attractive alternative production platform for the large-scale manufacture of valuable human drugs based on the mammary gland’s high protein production capacity, easy access to the recombinant proteins in milk, great scalability and cost effectiveness (6). The concept for such an animal-based production system has already been fully validated with the approval of three proteins isolated from milk. Recombinant forms of human antithrombin produced in the milk of goats, as well as C1 esterase inhibitor and coagulation factor VII produced in the milk of rabbit,s are all being commercialized as therapeutics for the treatment of human diseases (7-11). Moreover, the potential for mass production of a therapeutic antibody was exemplified in a recent study reporting production of an anti-CD20 antibody in the milk of cattle (12).

Cetuximab is a chimeric mouse-human mAb, targeting the epidermal growth factor receptor (EGFR), that has approval for the treatment of metastatic colorectal and head and neck cancers (13). The approved mAb is produced in the mouse-derived cell line SP2/0 and is marketed under the brand name Erbitux. In 2014 it was the 19th biggest-selling drug worldwide, with annual sales of US$1.9 billion (1). However, the murine SP2/0 production cell line of commercial Erbitux modifies the variable region of the heavy chain with a galactose-α-1,3-galactose (α-Gal) at an N-linked glycosylation (14, 15). This glycan structure is not found in humans and has been shown to be associated with adverse effects in patients treated with Erbitux (14).

Here, we describe the development and characterisation of two transgenic goat lines to produce cetuximab in milk. We demonstrated trans-generational stability of the integrated transgenes and cetuximab production at levels up to 15 g/L of milk. gCetuximab was lacking the problematic α-Gal glycosylation and showed enhanced antibody-dependent cell-dependent cytotoxicity (ADCC) compared to commercial Erbitux, both hallmarks for the production of an improved cetuximab version.

## MATERIALS AND METHODS

All experiments were performed in accordance with the relevant guidelines and regulations with approvals from New Zealand’s Environmental Protection Authority and the Ruakura Animal Ethics Committee.

### Generation of Expression Vectors

Cetuximab light and heavy chain coding sequences were optimized and synthesized by Geneart AG, now a part of Thermo Fisher Scientific of Waltham, MA, USA. Two separate expression constructs were made to target expression of both the light and heavy chains of cetuximab to the lactating mammary gland. Both constructs were based on the expression vector Bc450, which contains an expression cassette consisting of two copies of a fragment of DNA isolated from the 5’ end of the chicken B-globin locus, which is predicted to reduce positional effects by directionally inducing transcriptionally active decondensed regions in chromatin (16), 5’ (6.2 kb) and 3’ (7.1 kb) goat β-casein gene (g*CSN2*) sequences flanking an XhoI cloning site that direct expression to the mammary gland, in a Supercos backbone. To generate the vector Bc2553, the cetuximab light chain was cloned into the XhoI site of a derivative of the expression vector Bc450 (Figure S1A).

The heavy chain expression vector Bc2584 was constructed by ligating three fragments of DNA (Figure S1B,C). The first step was to generate an intermediate construct by inserting the heavy chain coding sequence into Bc350, which is identical to Bc450 described above except for the replacement of the downstream SalI site with a NotI restriction site. Following excision of the heavy chain expression fragment with SalI and NotI, it was ligated in a second step with a 1.5 kb SalI/NotI fragment containing the puromycin resistance gene (17) and a SalI/SalI Supercos vector backbone fragment in a three-way reaction.

### Cell culture and cell clone isolation

Female primary goat fetal fibroblasts (GFFs) were cultured in Dulbecco’s Modified Eagle’s Medium (DMEM)/F12 with non-essential amino acids, 10% FCS and human fibroblast growth factor-2 (5 ng/mL, Sigma-Aldrich, St Louis, MO, USA) at 37 °C. GFFs were co-transfected with 3 µg of a 16.4 kb SalI fragment excised from the expression construct for the cetuximab light chain (LC) and 1.7 µg of a 18.7 kb SalI fragment for the heavy chain (HC) by nucleofection using the Basic Nucleofector Kit for Primary Mammalian Fibroblasts with programme A-24 according to the manufacturer’s instructions (Lonza, Cologne, Germany). Cell clones were isolated after approximately 12 days of puromycin selection (1.5 µg/mL), initially transferred individually into 96-well plates before transfer to larger multi-well formats for further expansion.

### Somatic cell nuclear transfer (SCNT)

*In vivo* ovulated oocytes were surgically flushed from synchronised donors and enucleated under minimal ultraviolet light exposure following staining with 2 µg/mL Hoechst 33342 (Sigma-Aldrich, St. Louis, MO, USA) for 2 min. Prior to SCNT, transgenic donor cells were serum starved for 4-6 days and then re-stimulated with medium containing 10% FCS approximately 4 h before fusion. Individual donor cells were fused with enucleated oocytes and simultaneously activated by two direct current electric pulses (2 kV/cm for 10 µs each) in fusion buffer containing 50 µM calcium. Approximately 45 min following successful fusion, reconstructed embryos were given either a second electrical treatment as above or were exposed to 2.5 µM ionomycin for 1 min to further stimulate activation. For the next 3 h, the SCNT embryos were cultured in cycloheximide (5 µg/mL) and cytochalasin B (5 µg/mL) and then cultured overnight in AgResearch SOF media. Resulting 1- and 2-cell embryos were surgically transferred to the oviducts of recipients two days after oestrus, at a range of 5-11 embryos/recipient. Pregnant goats were identified by ultrasound 30 days after transfer and monitored regularly thereafter. Parturition was induced for a planned delivery five days before expected full term by administering a combination of prostaglandin and dexamethasone 36 h beforehand.

### Genotype analyses

DNA was isolated from cell and blood samples using a Nucleon BACC2 kit (Cytiva, Little Chalfont, UK). Cell clones were analysed by PCR amplification of LC- and HC-specific amplicons. Insertion of the LC-transgene was determined with PCR primers GL560 and GL561. The HC-specific fragment (449 bp) was amplified with primer pair GL556/GL573. Amplification of a g*CSN2* fragment (198 bp) with primer pair GL579/580 served as positive control. The PCR amplification for each gene sequence was performed with 50 ng of genomic DNA in a 20 µL reaction using a KAPA Taq PCR Kit (Sigma-Aldrich, St Louis, MO, USA) and the following cycle conditions: one pre-cycle step 3 min at 95°C, 38 cycles of 15 s at 95°C, 15s at 60°C, 1 s at 72°C and a final step of 5 min at 72°C.

Confirmation of the endogenous to transgene sequence junctions were confirmed by PCR with primer pairs GL1313/GL1314 (BP1, GN388), GL1258/GL1256 (BP2, GN388) and GL1309/GL1312 (BP1, GN451). PCR conditions were as above except for a longer extension step of 5 s.

For ddPCR quantitation of cetuximab LC transgenes, 25 ng of genomic DNA were amplified with LC-specific primers GL1459/GL1460 and goat beta-lactoglobulin gene (*LGB*) primers GL1470/GL1471 as reference, in the presence of the LC target FAM probe GL1468 and reference HEX probe GL1472 in a 22 µL reaction. The probes were double-quenched with an internal ZEN quencher and 3’ IBFQ quencher (Integrated DNA Technologies, Coralville, IA, USA). Following droplet generation with a QX200 system (Bio-Rad), the DNA was denatured for 10 min at 95°C and amplified in each droplet by 40 cycles of 30 s at 94°C, 1 min at 60°C with a ramping rate of 2°C/s and a final step of 10 min at 98°C. Fluorescence of individual droplets was determined with the QX200 droplet reader and transgene copy number quantified using the QuantaSoft Analysis Pro software (Bio-Rad, Hercules, CA, USA). The copy number analysis was done with three technical replicates for each sample. The quantitation of the HC transgenes was performed the same way but used HC-specific primers (GL1457/GL1458) and probe (GL1467) instead of the LC-specific ones. All PCR primer and fluorescent hybridisation probe sequences are listed in Table 1. For Southern analyses, 5 µg of genomic DNA was digested with EcoRI, transferred to membrane and hybridized with a radioactively-labelled 660 bp XhoI/HindIII fragment, spanning parts of exon 7 and exon 8 of g*CSN2*, for the detection of LC- and HC-transgenes as well as the endogenous g*CSN2* fragments. Hybridization signals were determined by X-ray film densitometry (GS-800, Bio-Rad, Hercules, CA, USA) and transgene copy numbers quantified with normalization against the two endogenous g*CSN2* copies using Quantity One software (Bio-Rad, Hercules, CA, USA).

**Table 1.**
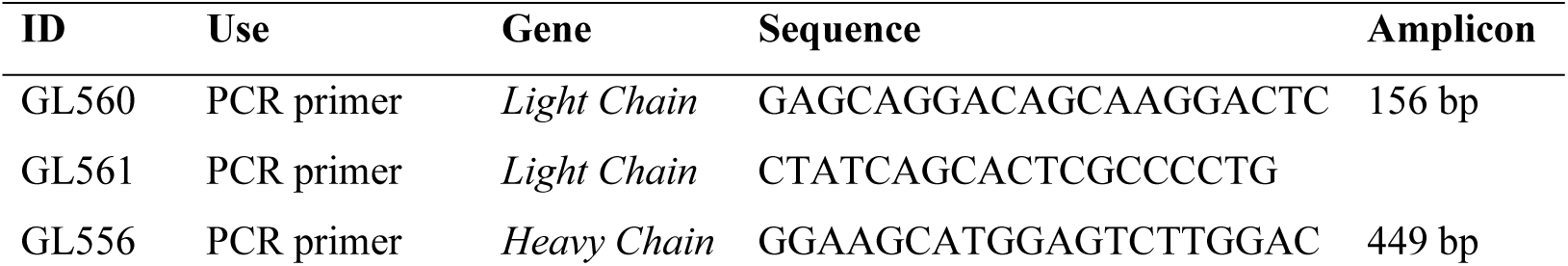
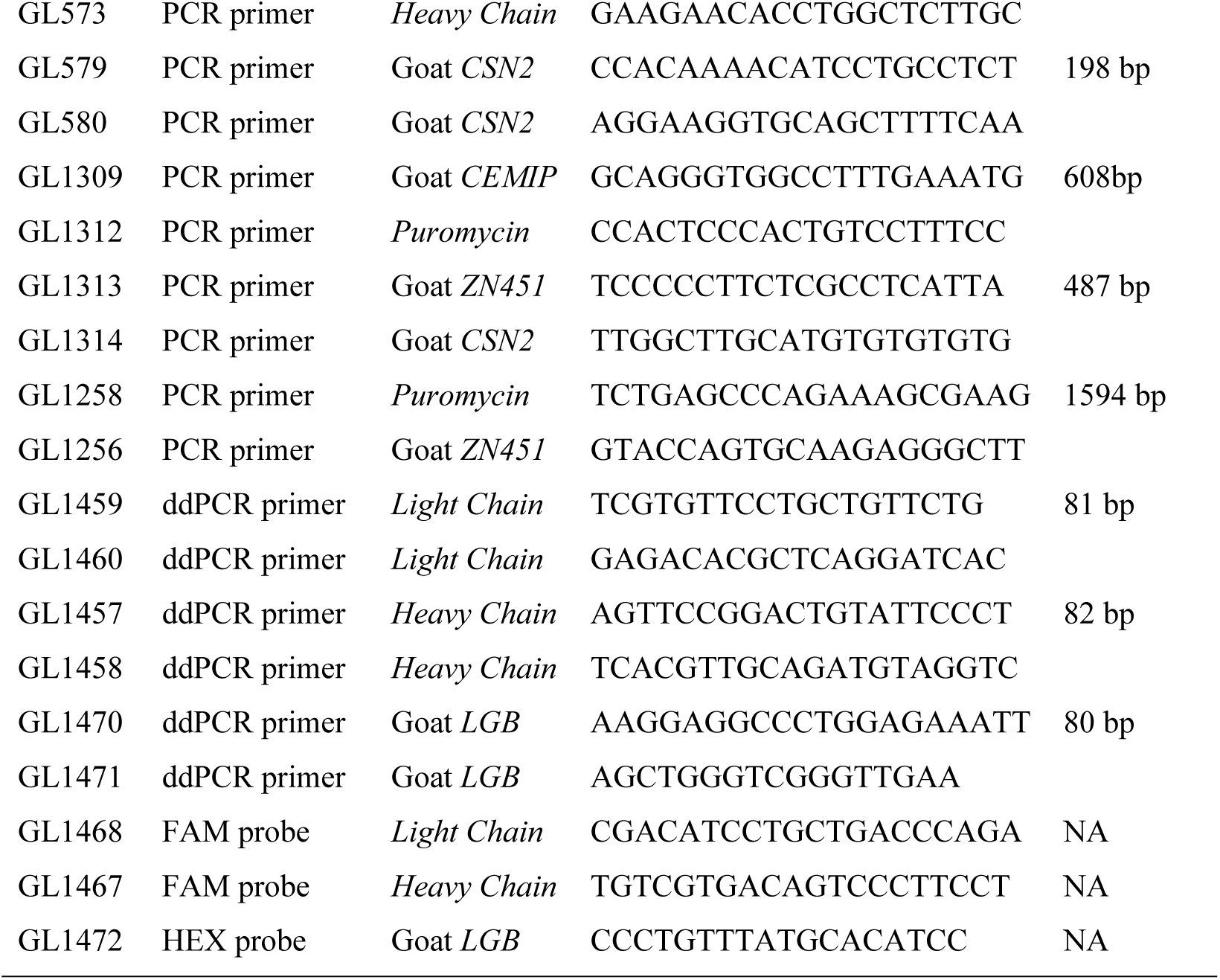
Summary of PCR primers and probes.

### Whole genome next generation sequencing

To locate and enumerate the transgenes, Illumina Next Generation sequencing was employed. Two libraries, Nextera mate pair (5kb) and Illumina PCR-free paired end (550bp), were made from a single GN388 and GN451 founder goat. The resulting fastq files were checked for low quality regions and adapters using fastqc version 0.11.5 (18). These regions were trimmed using trimmomatic version 0.3.6 (19). The trimmed reads were then mapped to the combined sequences of the two transgenes and Caprine Genome Assembly ASM170441 version 1 (20), using the “MEM” algorithm of BWA version 0.7.9a-r786 (21) with the resulting output being converted into BAM file and indexed using samtools version 1.3 (22). To locate the transgenes, the resulting output was searched for read pairs where each read mapped to a different chromosome or one read mapped to one of the transgene sequences and the other mapped to a chromosome. A single distinct location appeared for each of the two sequenced goats. This was checked and confirmed using the integrative genome viewer (23). The copy numbers of both transgenes were enumerated by comparing the average coverage of g*CSN2* and the average coverage of all and specific regions of the genome as calculated from the output of the “depth” function of samtools (22) selecting only sequenced bases with a quality value of at least 30 from sequencing reads with a mapping quality of at least 30. To enumerate copies of the LC- and HC-transgenes and ensure their integrity, the mapping of the reads across the unique regions encoding cetuximab light and heavy chains and not shared between transgene or endogenous host genome, was enumerated using the “depth” function of samtools and examined in Integrative Genomic Viewer for possible mutations. The overall copy number for both transgenes was enumerated by comparing the average coverage of the 5’ and 3’ g*CSN2* regulatory sequences (CM004567.1, position 86,000,517-86,007,945 and 86,013,215-86,019,039, respectively), shared by the genome and the transgenes, and the average coverage of all regions of the genome as calculated from the output of the “depth” function of samtools (22). In addition, we also determined the coverage across the CNS2 region which encompassed the coding information unique to the genome and not part of the transgenes (CM004567.1, position 86,007,946-86,013,214). The coverage per haploid genome was calculated by dividing the average coverage of all regions of the genome by two. To calculate the copy number of the transgenes and the endogenous g*CSN2* genes, the average coverage of the sequencing reads over the relevant regions was calculated and divided by the coverage per haploid genome. To enumerate copies of the LC- and HC-transgenes we used coverage for the transgene specific regions encoding the LC- and HC- chain which is not shared between transgene or endogenous host genome for the calculation. To ensure the integrity of the transgenes, the mapping of the sequence reads across the unique LC- and HC-regions was examined using Integrative Genomic Viewer for possible mutations and gaps in coverage.

### Induced lactation and milk analysis

At the age of 3-6 months, goats were hormonally induced into lactation as described previously (24). Briefly, every other day goats were given intramuscular injections of estrogen (0.25 mg/kg live weight) and progesterone (0.75 mg/kg live weight) for a total of seven treatments. For the following three days, goats received daily intramuscular injections of dexamethasone (0.3 mg/kg live weight). From the third estrogen/progesterone injection onwards, the mammary glands were manually massaged daily to stimulate lactogenesis and milk secretion was expected to begin at the end of the estrogen/progesterone treatment or with the dexamethasone injections. Once milk secretion had started, the goats were hand milked once a day and milk collected for up to 30 days.

For access to natural lactation milk, goats were either naturally mated or artificially inseminated. Following kidding, goats were milked twice daily with am and pm milk pooled to generate daily milk samples for analysis.

Milk proteins were separated by SDS PAGE on Criterion TGX 4-20% gels (Bio-Rad). Following wet transfer onto a Hybond ECL membrane (Cytiva, Little Chalfont, UK), expressed gCetuximab light- and heavy-chains were detected with a peroxidase-conjugated anti-human IgG antibody (MP Biomedicals, Santa Ana, USA) and visualized with Bio-Rad Clarity™ Western ECL Substrate (Bio-Rad, Hercules, CA, USA). Chemiluminescence was detected with an ImageQuant Las 4000 instrument (Cytiva, Little Chalfont, UK) and quantified using Bio-Rad Quantity One software (Bio-Rad, Hercules, CA, USA). For a reference standard we used an in-house produced monoclonal antibody of known concentration.

### Quantitation of Induced and Natural Lactation Milk Samples

Milk proteins were separated by SDS-PAGE on NuPage 4-12% Bis-Tris gels (Invitrogen, Carlsbad, CA, USA) electrophoresed in MOPS buffer and transferred to nitrocellulose membranes (Whatman, Little Chalfont, UK) in NuPage transfer buffer. Expressed cetuximab light- and heavy-chains were detected with an IR800-affinity purified goat anti-human IgG (H+L) (Rockland 609-132-123, Rockland Immunochemicals, Limerick, PA, USA) in Licor Blocking buffer and visualized on a Licor Odyssey Imaging System. Estimates of cetuximab concentration were made by visual comparison to standard antibody of known concentration. Estimates of concentration were also made by comparison of the yield of purified antibody to the starting volume of milk, adjusted for normal losses in purification.

### Purification of gCetuximab

Frozen milk samples were defrosted, and volumes of 25 mL and 100 mL of transgenic goat milk were processed for purification of gCetuximab. Milk samples were diluted 1:1 with PBS followed by centrifugation at 10,000 to 12,000 x g for 60 min at 4°C. The resulting clarified milk fraction was filtered through a 2.7 µm or 5 µm depth filter and stored frozen prior to commencing with the isolation of gCetuximab by column chromatography. For this next step, the clarified milk was thawed and loaded at 150 cm/h and 90 cm/h onto a 22 mL (1.6 x 11 cm) and 40 mL (1.6 x 20 cm) rPA Sepharose FF (GE Healthcare, Uppsala, Sweden) Protein A affinity chromatography column equilibrated with PBS for samples with a 25 mL and 100 mL milk starting volume, respectively. The column was washed with PBS followed by 100 mM acetate pH5.5 buffer containing 500 mM sodium chloride and gCetuximab eluted with 100 mM glycine pH3.0 buffer. The eluted fraction was neutralized with 2 M Tris pH7.0 and either diluted first 1:1 with 20 mM phosphate pH7.0 buffer (25 mL milk sample) or loaded directly (100 mL milk sample) onto a Q Sepharose FF (GE Healthcare, Uppsala, Sweden) equilibrated with 20 mM phosphate pH7.2 buffer column at 150 cm/h and 90 cm/h, respectively. gCetuximab was collected in the flow-through fraction and diafiltered against 5 diavolumes of PBS using 0.04 m^2^ of 30 kDa MWCO PES membrane (Novasep, Shrewsbury, MA, USA). Samples were quantified spectrophotometrically by absorbance at 280 nm using a general extinction coefficient of 1.4.

### gCetuximab glycosylation analysis

Milk samples were obtained and the gCetuximab purified as described above. Samples were analyzed by contract with Glycan Connections LLC (Lee, NH, USA) using LC/MS methodology to determine the glycosylation patterns. The percent of each of the resulting patterns were tabulated.

Relative galactosylation refers to the sum of the percentage of each carbohydrate structure found in the sample multiplied by the number of Gal residues in that structure, which can be zero, one, or two. For example, if a fully sialated complex structure (2 gals) were present at 15% of the carbohydrate structure population, it’s contribution to relative galactosylation would be 2 x 15 = 30. A similar hybrid structure with one Gal would contribute 1 x 15 = 15 to the relative galactosylation. By this method the maximum relative galactosylation could be 2 x 100% or 200.

Dividing this value by 2 gives the percentage of the maximum potential galactosylation in the sample.

Any given molecule contains four N-linked carbohydrate structures - two on the heavy chain constant regions and two on the heavy chain variable regions. Additionally, each carbohydrate structure can have one or two Gal residues, so the maximum number of Gal per molecule is eight. Multiplying the percentage galactosylation times eight gives the average number of Gal/mole of the sample.

### Western blot to determine presence of alpha 1,3 galactose

Western blot analysis was carried out using standard procedures for SDS-PAGE electrophoresis on 4-12% NuPage gels in MOPS buffer (Invitrogen, Carlsbad, CA, USA), then transferred to nitrocellulose membranes (Whatman, Little Chalfont, UK) in NuPage Transfer buffer. The detecting antibody used was a mouse IgM to alpha 1,3 gal (ALX-801- 090, Enzo Life Sciences, Farmingdale, NY, USA). The secondary antibody was HRP goat anti-mouse (SC2064, Santa Cruz Biotechnology, Dallas, TX, USA), followed by visualization with ECL Western Blot Detection Reagent (Amersham, Little Chalfont, UK) on X-ray film.

HPLC assay for α-Gal was performed by treating 50mg of protein in 50mM Sodium Acetate pH5.0 with and without α-galactosidase A (Fabrazyme, Sanofi Genzyme, Cambridge, MA, USA). The released α-Gal was collected in a Millipore 5K MWCO spin filter and labelled with 2-aminobenzoic acid (2AA), then analyzed by HPLC on a Waters Symmetry C18 column.

### Binding to EGFR

Binding to EGFR was evaluated with an assay to show binding to HTB-30 cells, a human breast cancer cell line expressing EGFR. Briefly, cells were washed with PBS and counted. 1×10^6^ cells were aliquoted and mixed with antibody samples at various concentrations, then incubated on ice for 30 min. Cells were again washed with PBS to remove unbound primary antibody and incubated with goat anti-human IgG conjugated to FITC (Southern Biotechnology Associates, Birmingham, AL, USA) for 30 min on ice. Cells were again washed and fixed with 1% paraformaldehyde, diluted to a concentration of 1×10^5^ cells/mL, and fluorescence determined on a Guava flow cytometer (Merck Millipore, Darmstadt, Germany). The mean fluorescence intensity (MFI) was plotted against antibody concentration to show binding efficiency.

### Binding to CD16

Binding to CD16 was studied by a competitive assay with an anti-CD16-PE/3G8 antibody conjugated to phycoerythrin (Beckman Coulter, Brea, CA, USA) and unlabelled test and control antibodies using Jurkat-CD16 cells.

In V bottom 96 well plates, 2×10^5^ cells/well (50 µL of cells at 4×10^6^ cell/mL) were incubated 20 min at 4°C with a final concentration of 333 µg/mL of the tested molecules diluted in PBS, simultaneously with anti-CD16-PE/3G8 conjugate diluted at 1/10, a fixed concentration. Cells were washed by adding 100 µL PBS and centrifuged at 1700 rpm for 3 min at 4°C. Supernatant was removed and 300 µL cold PBS was added. Binding of anti-CD16-PE/3G8 to CD16 expressed by Jurkat-CD16 cells was evaluated by flow cytometry. Relative CD16 binding is expressed in percent; 100% being the MFI obtained with the R297 anti-D mAb (LFB, Les Ulis, France) and 0% the MFI measured in the presence of Rituxan (commercial). Antibody R297, produced in YB2/0 cells, is used as an example of a low fucose antibody with high affinity for CD16, while Rituxan, produced in CHO cells, has significantly lower affinity for CD16, providing high and low binding examples used as arbitrary 100% and 0% standards. (Data not shown.)

### Presence of EGFR on Hep-2 Cells for Use in Activation Assay

EGFR expression in Hep-2 cells was confirmed by flow cytometry. Briefly, Hep-2 cells and antibodies were diluted in diluent (PBS with 1% FCS) and 1 x 10^5^ cells were incubated with 100 µL of Erbitux at 10 and 50 µg/mL (final concentration) at 4°C for 30 min.

After washing in diluent, bound antibodies were visualized using 100 µL of a 1:100 dilution of goat F(ab’)2 anti-human IgG coupled to phycoerythrin (Beckman Coulter, Brea, CA, USA) at 4°C for 30 min. The cells were washed and MFI was quantified by flow cytometry (FC500, Beckman Coulter, Brea, CA, USA).

### Jurkat-CD16 activation assay

Hep-2 target cells were incubated with increasing concentrations of anti-EGFR antibodies (0 to 500 ng/mL) in the presence of Jurkat CD16 cells and the protein kinase C activator phorbol-myristate acetate (P8139, Sigma-Aldrich, St Louis, MO, USA). After 16 h of incubation at 37°C, the quantity of IL-2 released by Jurkat-CD16 cells was measured in the cell supernatant by colorimetry using a human IL-2 DuoSet ELISA DY202-05 Kit (R&D Systems, Minneapolis, MN, USA).

## RESULTS

### Generation of gCetuximab-producing goats

Female goat fetal fibroblast (GFF) primary cell lines were co-transfected with two separate constructs for the mammary gland-specific expression of the light chain (LC) and heavy chain (HC) of cetuximab (Figure 1A).

**Fig. 1.**
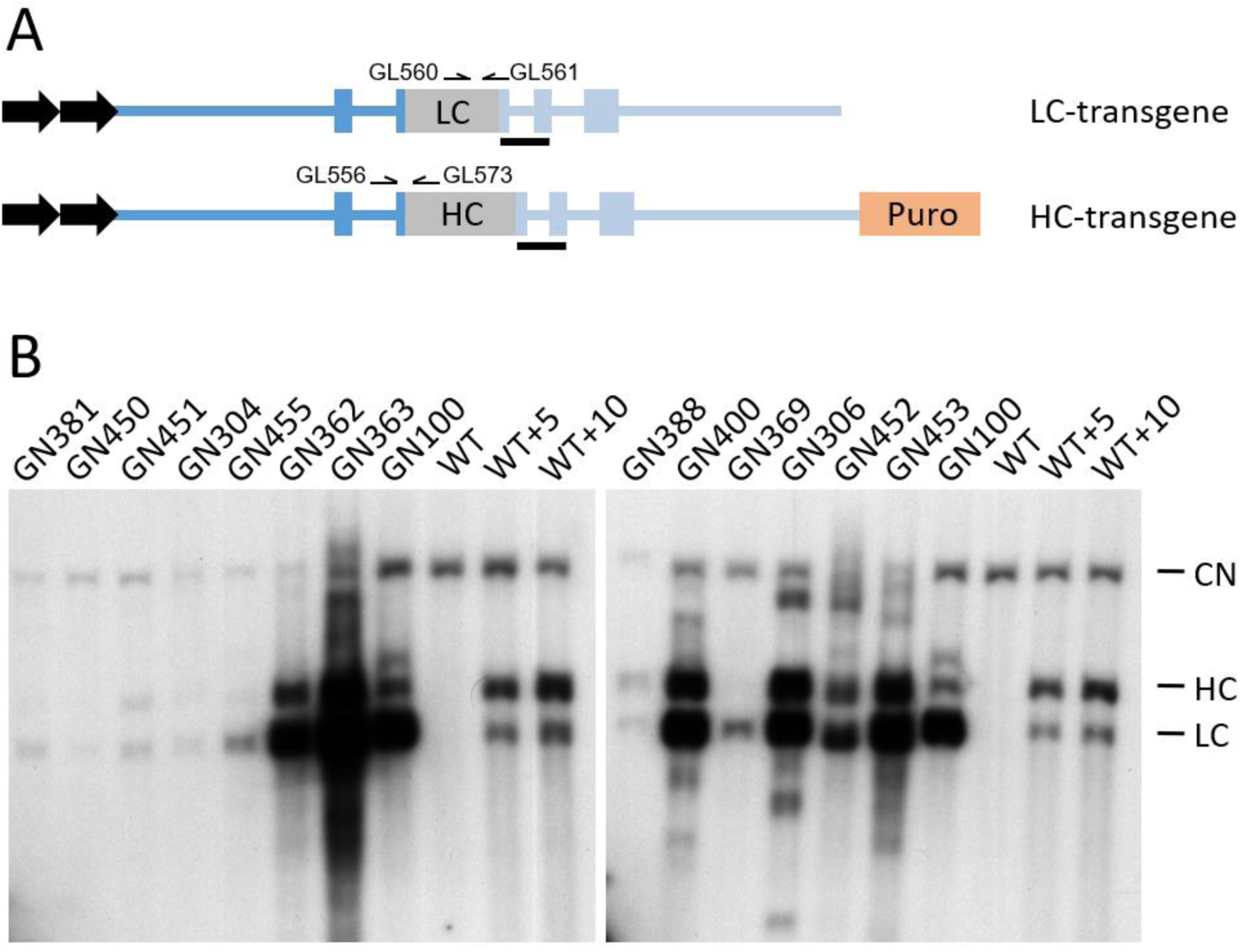
gCetuximab expression construct outline and detection. A) Schematic of the transgenes encoding the cetuximab light chain (LC) and heavy chain (HC) under control of the 5’ (dark blue) and 3’ (light blue) g*CSN2* regulatory sequences used for the co-transfection of primary fetal goat cells. The blue boxes depict exons 1, 8, 9 and partial exons 2 and 7 of g*CSN2* that flank the LC and HC coding regions. The transgenes also included tandem chicken β-globin insulators (black arrows) and at the 3’ end of the HC-transgene an antibiotic selection marker (Puro) driven by the 3-phosphoglycerate kinase (PGK) promoter. The position of primers used for PCR genotyping and the Southern hybrydisation probe (black bar) are indicated. B) Southern analysis of transgenic cell clones. Depicted are the resulting hybridisation signals of genomic DNA from a selection of different cell clones probed with a g*CSN2* fragment. CN: signal from the endogenous g*CSN2* copies; HC: signal of the heavy chain encoding transgene; LC: signal of the light chain encoding transgene; WT: non-transgenic wild type control; WT+5/10: wild type genomic DNA spiked with the equivalent of 5/10 transgene copies.

From two independent transfection campaigns performed within the laboratory over two distinct time periods, we isolated a total of 23 cell clones that were confirmed with stable integration for both of the transgenes identified by PCR. The number of LC- and HC-transgene copies was determined by Southern analysis, revealing a wide range in copy numbers across the individual cell clones (Figure 1B, Table S1).

We selected nine primary cell clones, representing low, medium and high transgene copy numbers that were used as donor cells to generate founder goats by somatic cell nuclear transfer (SCNT). Transgenic female kids were born from all nine cell clones. Overall efficiency for the production of live kids per total number of transferred embryos for the different cell clones ranged between 1% and 8% (on a per embryo transfer basis of calculation) to term as well as to weaning (Table 2).

**Table 2.** Summary of nuclear transfer results.

| **TG Line** | **Embryos transferred**<br>**(# of recipients)<sup>a</sup>** | **Day 30 of gestation**<br>**(# of recipients)<sup>b</sup>** | **Day 100 of gestation**<br>**(# of recipients)<sup>b</sup>** | **Development**<br>**to term (%)<sup>c</sup>** | **Development**<br>**to weaning (%)<sup>c</sup>** |
| --- | --- | --- | --- | --- | --- |
| **GL6.7** | 56 (8) | 2 (2) | 2 (2) | 2 (4) | 2 (4) |
| **GL6.7** | 57 (10) | 11 (5) | 5 (3) | 2 (4) | 1 (2) |
| **GL8.3** | 54 (10) | 5 (3) | 4 (3) | 4 (7) | 4 (7) |
| **GN97** | 25 (5) | 1 (1) | 0 (0) | 0 (0) | 0 (0) |
| **GN97** | 48 (7) | 1 (1) | 2 (1) | 2 (4) | 1 (2) |
| **GN97** | 45 (8) | 3 (3) | 3 (2) | 2 (4) | 2 (4) |
| **GN99** | 26 (6) | 3 (2) | 2 (1) | 2 (8) | 2 (8) |
| **GN100** | 77 (10) | 5 (4) | 5 (4) | 4 (5) | 3 (4) |
| **GN118** | 100 (12) | 1 (1) | 1 (1) | 1 (1) | 1 (1) |
| **GN304** | 24 (5) | 1 (1) *Day 49 | 1 (1) | 1 (4) | 1 (4) |
| **GN388** | 25 (4) | 1 (1) | 0 (0) | 0 (0) | 0 (0) |
| **GN388** | 35 (7) | 2 (2) *Day 52 | 2 (2) | 2 (6) | 2 (6) |
| **GN451** | 37 (7) | 4 (3) | 4 (3) | 3 (8) | 1 (3) |

<sup>a</sup> Total number of 1- 2-cell embryos transferred to the total number of recipients given in brackets.
<sup>b</sup> Total number of fetuses detected by ultrasound scanning at the indicated time of gestation (\* alternative gestational age examined). The values in brackets indicate the number of pregnant recipients.
<sup>c</sup> Total number of live goats at term or weaning (approximately 3 months) and percentage of total number of transferred embryos that develop into live goats in brackets.

Most of the kids were born as singletons (17 kids) while four sets of twins were also produced. Although most kids were strong and healthy after birth, five kids died within the first five days. This included one singleton that suffered complications from milk bloat and another singleton as well as three twin kids that were relatively small and weak when they were born. PCR analysis was used to confirm the genotype of all founder kids which demonstrated the presence of the light chain and heavy chain transgenes (Figure S2).

To determine whether the different transgenic lines expressed gCetuximab in their milk, the founders for the nine transgenic lines were hormonally induced into lactation and milked for up to 30 days. The two transgenic lines with very high numbers for both LC and HC transgene copies (GL8.3 and GN118) failed to lactate. The cloned founder goats from the GL8.3 line only secreted a single drop of a highly viscous substance. Similarly, the cloned founder goats from the GN118 line only produced secretions over three days that ranged from a maximum of 1 mL to a few drops of viscous material and failed to properly lactate. These lines were not further considered for gCetuximab production. The three transgenic lines GN97, GN99 and GN100 with medium (GN97) to high (GN99, GN100) LC transgene copy numbers and low HC transgene copy numbers produced milk secretions for at least nine days (Figure 2A). However, they were unable to maintain the lactation over time and the milk yields dropped over a relatively short period, until milk secretion stopped after nine to 25 days, depending on the transgenic line and individual animal. In contrast, low transgene copy lines GN304, GN451 and GN388 maintained lactation and increased in yield with progression of lactation. The milk production levels were markedly higher for both founder goats from line GN388 (maximum of 270 mL and 100 mL per day, respectively) compared to lines GN304 and GN451 (maximum of 23 mL and 22 mL per day, respectively). Founders from the GL6.7 line that were hormonally induced to come into lactation were only milked for one day to confirm expression and not assessed for milk yields over several weeks.

**Fig. 2.**
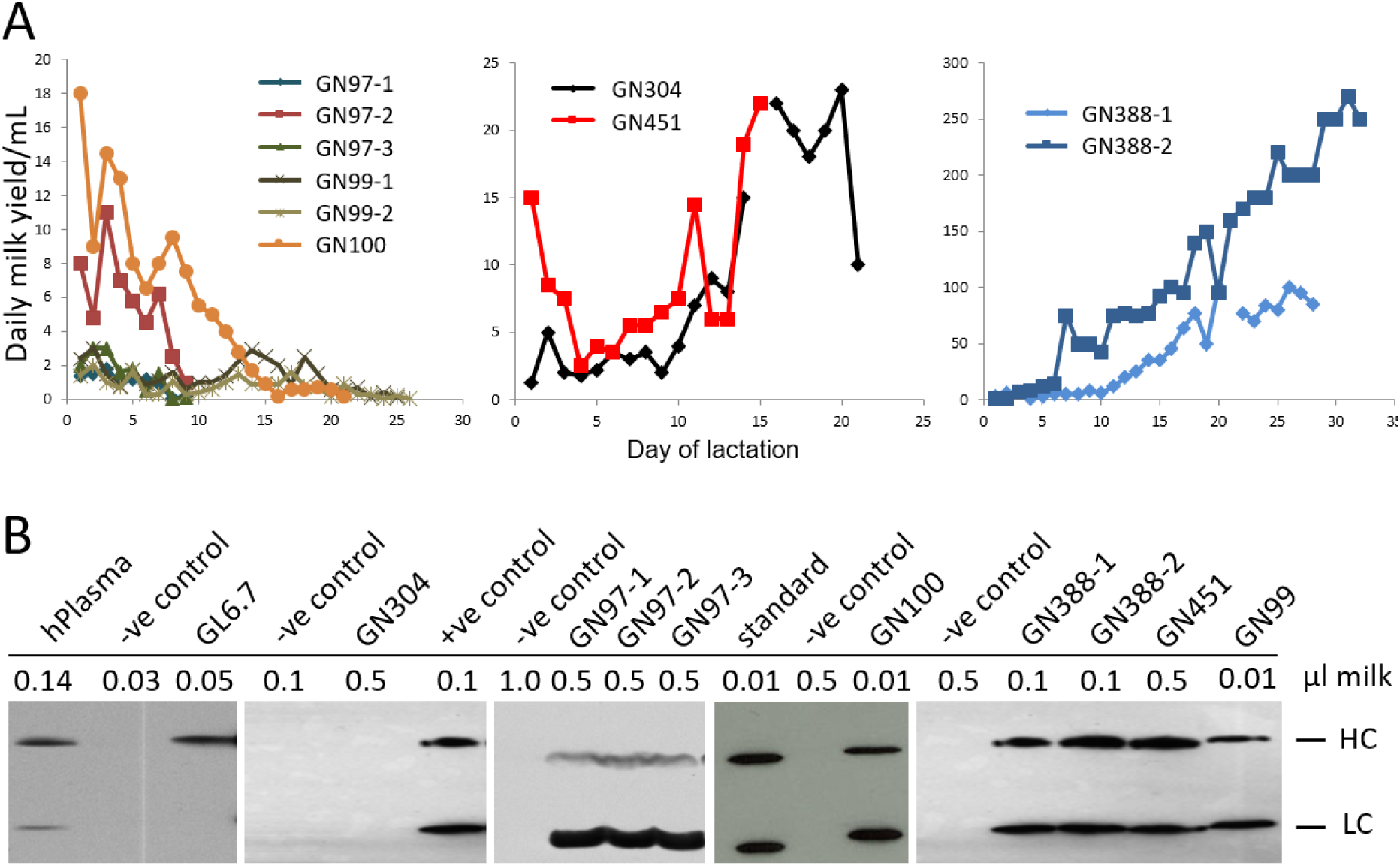
Milk and gCetuximab production by founder goats from a hormonally induced lactation. A) Daily milk yields for individual founder goats for the medium copy number transgenic lines GN97, GN99 and GN100 (left panel), for the founder goats of the low transgene copy number lines GN304 and GN451 (center panel) and founders of the low transgene copy number line GN388 (right panel). B) gCetuximab expression in milk. Western results of induced milk samples, produced by goats of the indicated transgenic lines, probed with an anti-human IgG antibody detecting gCetuximab light-(LC) and heavy-chain (HC). hPlasma: milk sample spiked with human plasma; -ve control: milk sample, not containing gCetuximab; +ve control: milk sample known to contain gCetuximab; standard: monoclonal antibody at 15 µg/µL.

Milk samples were analyzed by western blotting to detect and determine the level of gCetuximab expression. This analysis revealed that the founder transgenic line GL6.7 only expressed the gCetuximab heavy chain whereas line GN304 failed to express either of the gCetuximab chains (Figure 2B). The remaining five founder transgenic lines (GN97, GN99, GN100, GN388, GN451) expressed both transgenes with gCetuximab expression levels ranging from approximately 5 g/L to 15 g/L (Figure 2B, Table 3). Based on the hormonal induced lactation profiles and gCetuximab expression levels, two primary founder transgenic lines were selected (GN388 and GN451) for further characterization and moving forward into developmental use in subsequent experiments.

**Table 3.** gCetuximab expression levels in different transgenic goat lines.

| **Line** | **gCetuximab expression in milk** |
| --- | --- |
| GL6.7 | Good HC expression but no expression of LC |
| GL8.3 | Failed to lactate |
| GN97 | Unequal expression (low HC, high LC) |
| GN99 | ~ 15 g/L |
| GN100 | ~ 15 g/L |
| GN118 | Failed to lactate |
| GN304 | No detectable levels of HC and LC |
| GN388 | ~ 5-10 g/L |
| GN451 | ~ 3-5 g/L |

### Chromosomal transgene integration in lines GN388 and GN451

The transgene integration sites for both lines taken forward in this study were determined by whole genome next generation sequencing which revealed a single distinct chromosomal location for each transgenic line (Figures S3A-D). In line GN388, the transgenes were integrated into a single insertion site on chromosome 23 at position 45,650,975 (CM004584.1) by replacing 22 bp of endogenous sequence (45,650,975-45,650,997). For line GN451 the transgene insertion site was found on chromosome 21 at position 26,329,477 (CM004584.1). Mapping of the endogenous sequences flanking the breakpoints to the goat genome determined that in line GN388 and line GN451 the transgenes were inserted within the last intron of the longest isoform of the zinc finger protein 451 gene (*ZNF451*) and the first intron of the cell migration inducing hyaluronan binding protein gene (*CEMIP*), respectively. The breakpoints were verified by PCR and sequencing of the PCR products (Figure 3). This confirmed both breakpoints in line GN388 with the 5’-end of a partial transgene copy at one side and the 3’-end of the HC-transgene at the other breakpoint. For line GN451, only one of the insertion breakpoints could be determined by PCR because of the presence of a repetitive sequence in the genomic region at the other side. At the identified breakpoint, the endogenous genomic sequence was juxtaposed to the 3’ terminal end of a transgene copy. PCR amplification between the endogenous locus and the puromycin selection marker verified the breakpoint and identified that the 3’ end of the HC-transgene was present at the breakpoint.

**Fig. 3.**
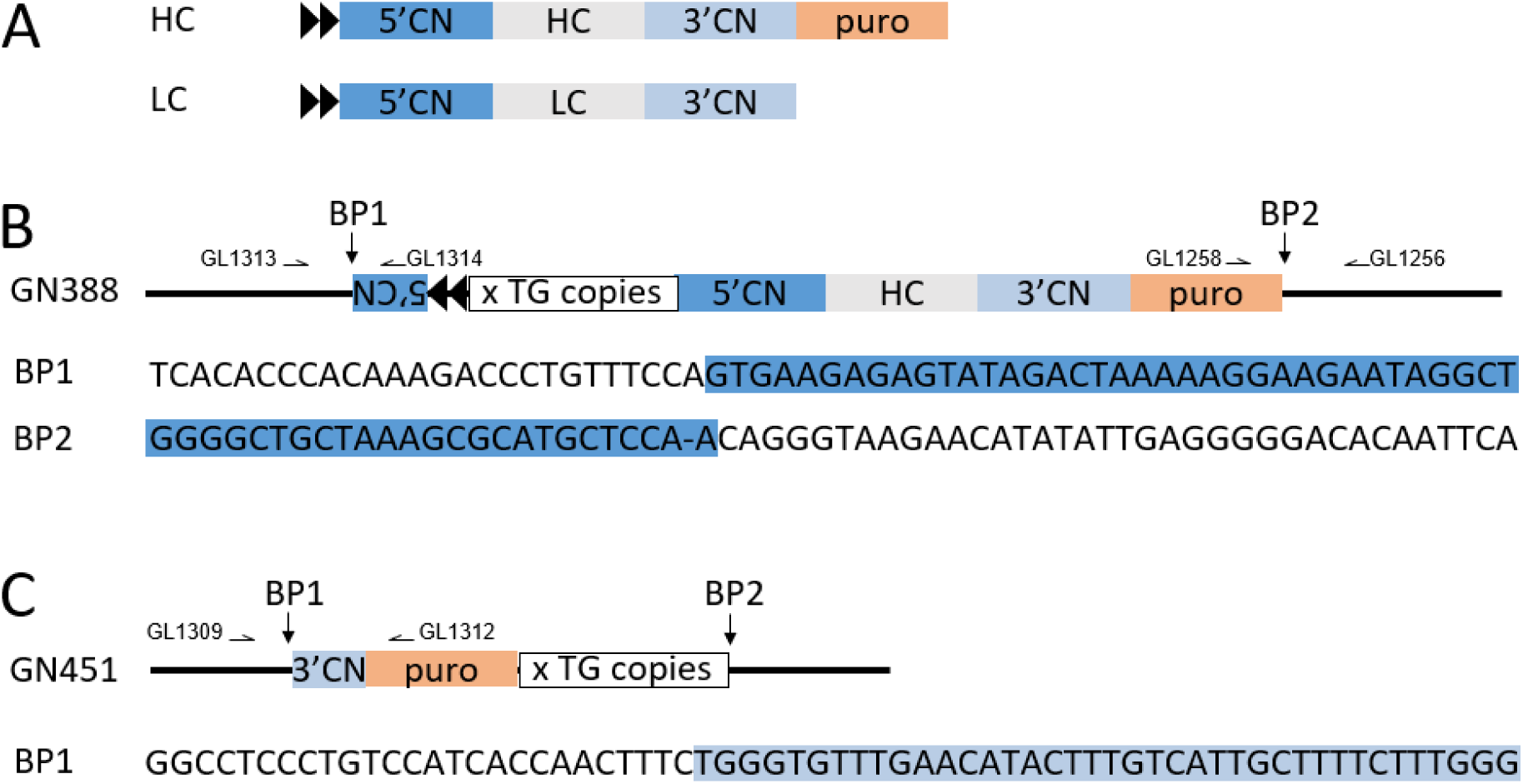
Transgene insertion sites. A) schematic representation of the functional elements of the LC- and HC-transgenes. 5’CN, 3’CN: g*CSN2* 5’ and 3’ regulatory sequences; HC, LC: heavy and light chain encoding sequences; puro: puromycin selection marker; black arrowheads: chicken β-globin insulators. B) schematic outline of the transgene insertions in line GN388. The transgenes at the breakpoints (BP1, BP2) of the endogenous locus are depicted. The corresponding endogenous (black) and transgene sequences (highlighted in dark blue) at the breakpoints are shown below the insertion site outline. x TG copies: cluster of additional transgene copies. C) schematic outline of the transgene insertions in line GN451. The sequence of the junction between endogenous locus and transgene sequences (highlighted in blue) are shown for BP1. Location of PCR primers for amplification of the breakpoint junctions are indicated.

The next generation sequence data were also used to enumerate the number of transgene copies. From this analysis, we determined that line GN388 carried a total of seven to eight transgene copies comprised of two LC- and five HC-transgenes based on the mate pair library sequence data (Table S2A). Results with the paired end library were less consistent with 10-11 transgenes in total, comprised of one LC- and three HC-transgene copies. Line GN451 had a total of two transgene copies consisting of one LC- and one HC-transgene (Table S2B).

Next, we used REF to visually verify the integrity of the coding regions of the LC- and HC-transgene copies. For the transgenes in line GN451, the minimum coverage was 2 reads with an average coverage of 5.2 (paired end) and 6.5 (mate pair) for the LC-transgene and 4.2 (paired end) and 9.2 (mate pair) for the HC-transgene. Only reads with a mapping quality of 30 and a base call quality of 30 were considered, verifying the integrity of both single copy genes. For line GN388, the minimum coverage for the LC and HC transgenes was 17 and 30 for the paired-end library and 7 and 12 for the mate pair library, respectively giving an expected minimum coverage of any transgene copy of 2 per library and 8 when both libraries were combined. Using the binomial distribution, this gives a 1 in 14 and 1 in 800 chance, respectively, of an individual transgene copy not being represented in the worst case. No mutations or insertions were detected and there was relatively even coverage along the transgenes in both sequencing library types, indicating the coding regions of all copies of both LC- and HC-transgenes were intact in both transgenic lines (Figure S4).

### Stability of genotype

Stability of integration was demonstrated via transmission of the intact transgenes through to subsequent generations. PCR-amplification and sequencing of the breakpoints from the founders, F1 and F2 animals were performed for both transgenic lines. The amplified fragment size and sequences of the breakpoints in the offspring were identical to those of the founder animals for each of the respective transgenic lines (Figure S5).

Southern analysis (Figure 4A) and ddPCR assays (Figure 4B) were subsequently performed to determine whether the number of transgene copies remained stable or were subject to change in subsequent generations. Results from both methods were consistent and determined the same number of transgene copies for the two transgenic lines which were also in accordance with the sequence-based copy number results. For the founder goat from line GN388, we detected two light chain and five heavy chain transgene copies and one copy each of light chain and heavy chain for the founder goat from line GN451. F1 and F2 offspring from both founder transgenic lines, as well as F3 offspring from the transgenic line GN388, maintained the transgene copies present in the founders which confirmed stable inheritance and constancy of the transgenic genotype.

**Fig. 4.**
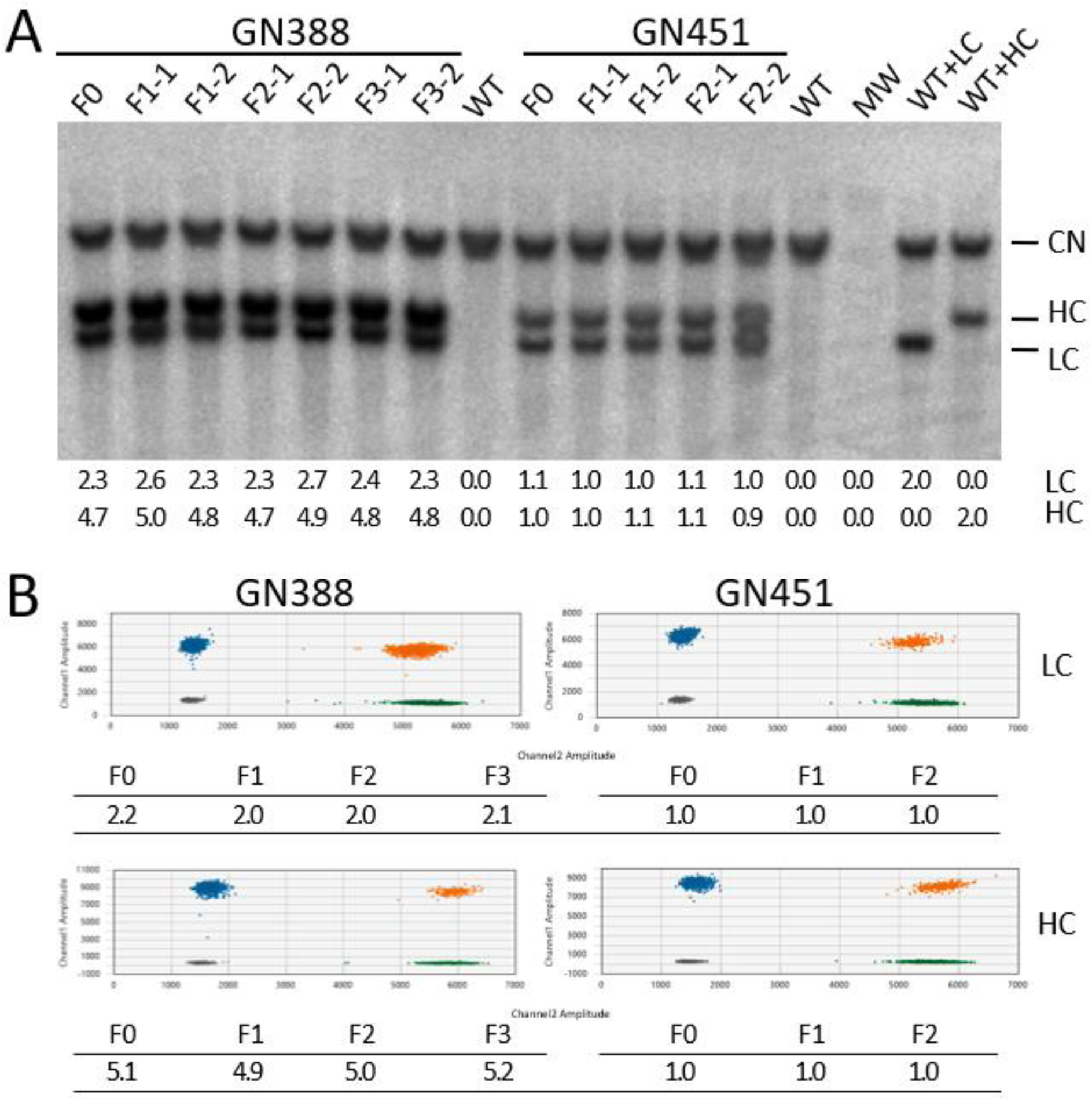
Transgene copy number determination in multiple generations. A) Assessment of transgene insertions by Southern analysis. Shown are the results for the founders (F0), two F1 (F1-1, F1-2), two F2 (F2-1, F2-2) and two F3 (F3-1, F3-2) offspring for the transgenic lines GN388 and GN451 as indicated. WT: non-transgenic wild type control; MW: molecular weight marker; WT+LC: wild type genomic DNA spiked with the equivalent of 2 LC copies; WT+HC: wild type genomic DNA spiked with the equivalent of 2 HC copies; CN: signal from the endogenous g*CSN2* copies; HC: signal of the heavy chain encoding transgene; LC: signal of the light chain encoding transgene. Quantitation results for the LC and HC transgene copies are summarized in the table below the Southern blot. B) Analysis by ddPCR. Shown are representative 2D-plots (channel 1 FAM-target vs channel 2 Hex-reference) for the ddPCR copy number variation assay results for the LC- and HC-transgenes in lines GN388 and GN451. Grey droplets: double negative (no DNA); blue droplets: LC/HC positive, green droplets: LGB reference positive; orange droplets: double positive for LC/HC and LGB. Tables below the 2D-plots summarize the calculated transgene copy numbers for the F0-F3 generations as indicated.

### gCetuximab expression levels in milk

To investigate consistency of the gCetuximab expression phenotype, we analyzed several goats. Daily lactation volumes of natural milk samples produced by F1 and F2 females from the transgenic line GN388 are presented in Fig. 5A, and F1 female offspring from the transgenic line GN451 (Fig.5B).

**Fig. 5.**
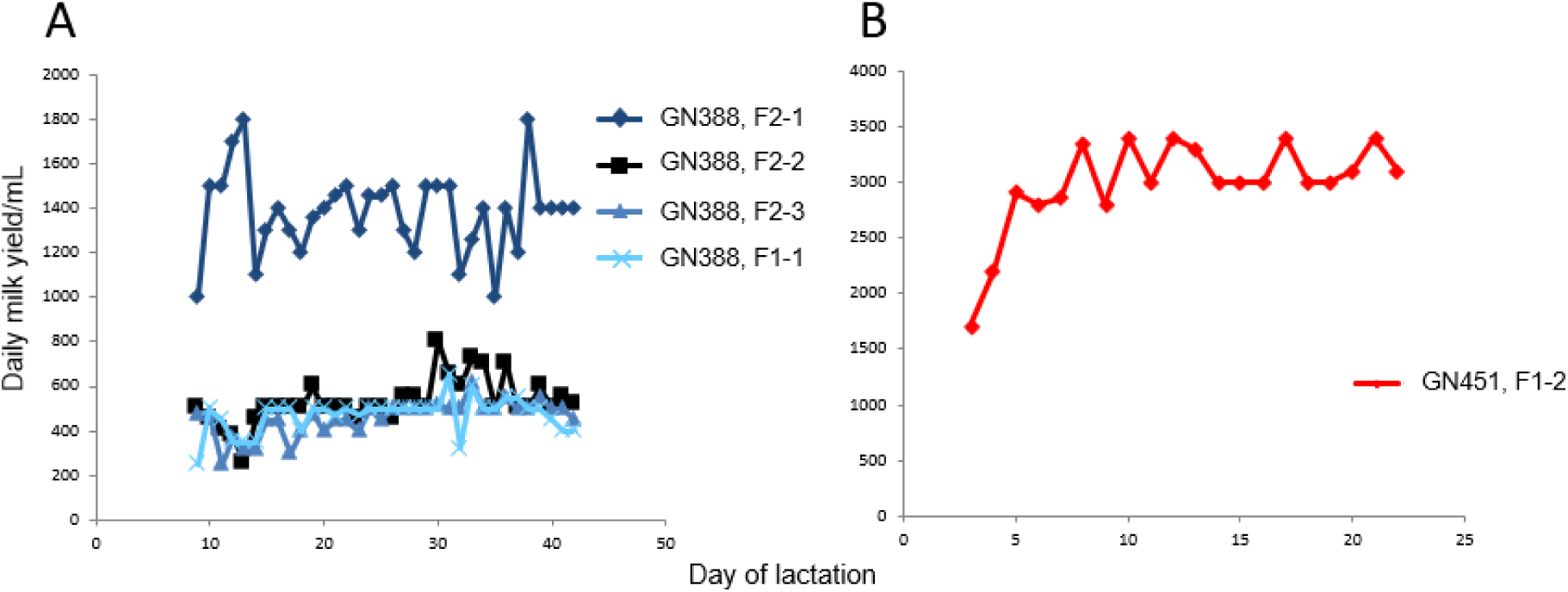
Milk yields of F1 and F2 generation goats. A) Daily natural lactation milk yields for different F1 and F2 goats of line GN388. B) Depicts natural daily milk yields for the F1-2 goat, line GN451. Please note the different scales for A and B.

The gCetuximab expression levels in milk were assessed by western blot and by calculation based on the gCetuximab yields determined according to our purification protocols. These estimates show similar gCetuximab production levels in the milk of the founders compared to their offspring with approximately 5-10 g/L for line GN388 and 3-5 g/L for line GN451 (Table 4).

**Table 4.** Estimated expression levels of goats from lines GN388 and GN451.

| Line | Goat | gCetuximab<br>(mg/mL) |
| --- | --- | --- |
| GN388 | F0 | 5<sup>a</sup>, 10<sup>b</sup> |
|  | F1-1 | 9<sup>b</sup> |
|  | F2-1 | 7<sup>b</sup> |
| GN451 | F0 | 5<sup>a</sup>, 3<sup>a</sup> |
|  | F1-1 | 3<sup>b</sup> |
|  | F1-2 | 3<sup>b</sup> |

<sup>a</sup> Estimate from western blot
<sup>b</sup> Estimate from purification based on yield and milk volumes.

### gCetuximab purification

gCetuximab was purified from milk samples produced by a founder goat of transgenic line GN100 (GN100, F0), and F1 goats for lines GN388 (GN388, F1) and GN451 (GN451, F1) using a two-step column chromatography strategy, comprising affinity and anion exchange separation principles. Purification from a 100 mL milk sample processed for line GN100 yielded a total of 850 mg gCetuximab. For lines GN451 and GN388, 25 mL of milk were processed for gCetuximab purification which yielded 55 mg and 150 mg purified antibody. The antibody preparations were then used for subsequent studies to further characterise the properties of gCetuximab.

### Glycosylation of gCetuximab

Glycosylation of cetuximab when produced in the goat mammary gland or in the mouse cell line will not be identical. Such differences could enhance the functionality of gCetuximab compared to Erbitux and provide evidence that the milk expession system could offer advantages to existing modalities of monoclonal antibody production.

Cetuximab carries an N-linked glycosylation site at aa 88, in the variable region of the heavy chain. The bi-antennary sugar moiety on this site has been found to carry α-Gal, when the commercial antibody Erbitux is produced in the Sp2/0 mouse cell line (25). This sugar modification is undesirable because it is highly immunogenic and has been shown to be responsible for adverse immune reactions following treatment with Erbitux (14). To investigate whether the antibody is modified by α-Gal when produced in the mammary gland of goats, purified gCetuximab derived from goat lines GN100, and GN388 was analyzed. Their glycosylation patterns were compared to commercially obtained Erbitux. Results shown in Figure 6 confirmed the presence of α-Gal in the commercial product and more importantly, absence of the α-Gal modification on the mAb produced in the milk of the transgenic goats.

**Fig. 6.**
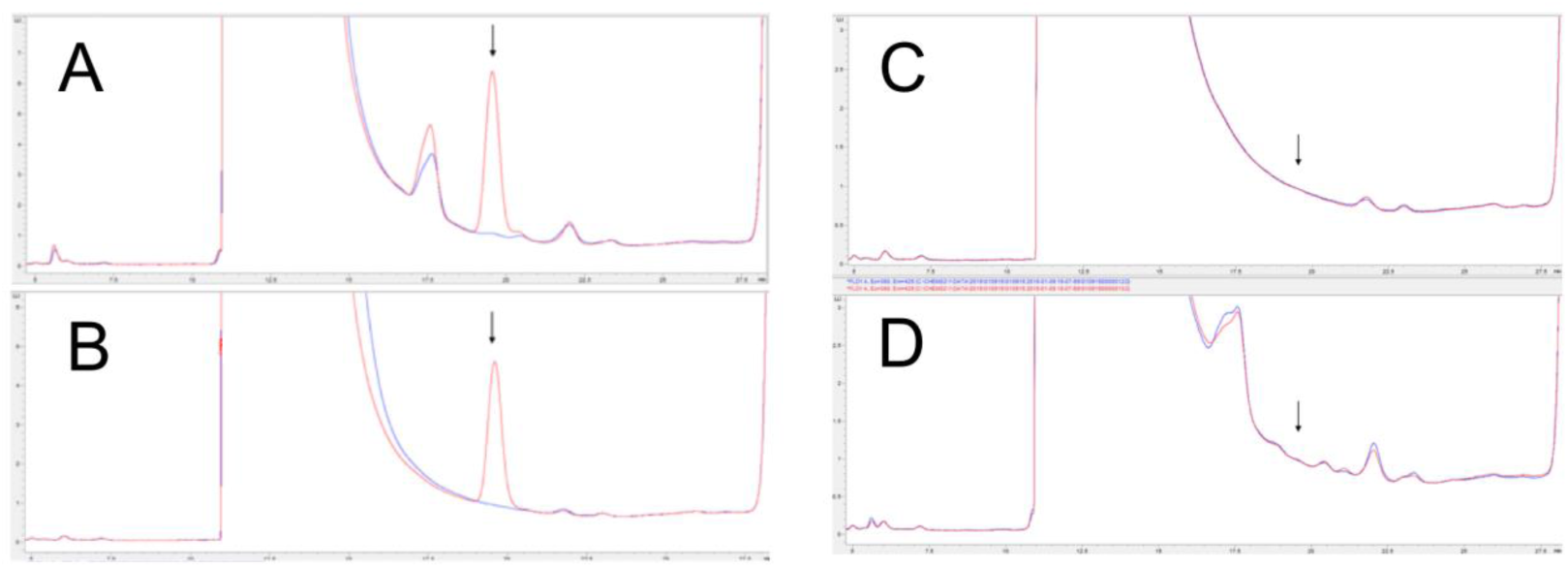
HPLC analysis of α -Gal on commercial and gCetuximab Chromatograms above show the release of α -Gal when treated with α -galactosidase A. Red traces show protein samples treated with α -galactosidase A, and blue traces are samples without added enzyme. Arrows indicate the position of the α -Gal peak, which elutes from the column at approximately 19.5 min. A) Laminin, a protein known to contain α -Gal; B) commercially sourced Erbitux; C) purified gCetuximab sample from the founder of line GN388; D) purified gCetuximab sample from the founder of line GN100.

In order to further confirm the lack of α-Gal on gCetuximab, western blot analysis was performed using a detecting antibody specific to the α-Gal moiety. Purified samples of the milk-produced gCetuximab and commerical Erbitux were tested for the presence of the α-Gal marker. Positive signals were detected on commercial Erbitux and goat serum (Figure S6). Since the goat adds α-Gal to serum proteins, it confirms the ability of the detecting antibody to identify the moiety. As can be seen, however, there are no signals indicating the presence of α-Gal in the purified gCentuximab (Figure S6). This corroborates the LC/MS results demonstrating that although the goat has the ability to add α-Gal and does so for serum proteins, goats do not decorate gCetuximab produced in the mammary gland and secreted into the milk.

In addition to the α-Gal modification, the gCetuximab samples were analyzed for sugar configurations. From the LC/MS data, the amounts of galactose (Gal) and afucosylated core sugars were calculated. As shown in the summary Table 5, the GN451 sample carried a higher level of afucosylated sugar, which has been correlated to increased CD16 binding and ADCC activity (26). In addition, increased galatose levels have also been correlated to increases in CD16 binding and ADCC activity, though to a lesser degree (27). This prompted us to further investigate CD16 binding and ADCC activity of gCetuximab. It should be noted that two of three goats from the GN388 line produced antibody with high levels of afucosylated antibody. Such variability has been observed and is attributed to variable genetics of the particular milking animal (data not shown).

**Table 5.** Summary of gCetuximab sugar configuration.

| **Antibody** | **Source** | **Relative galactosylation<sup>a</sup>** | **Gal/mole<sup>b</sup>** | **% afucosylated** |
| --- | --- | --- | --- | --- |
| gCetuximab | GN100, F0 | 185 (92%) | 7.4 | 8 |
| gCetuximab | GN388, F0 | 161 (81%) | 6.5 | 17 |
| gCetuximab | GN388, F1-1 | 151 (76%) | 6.0 | 6 |
| gCetuximab | GN388, F2-1 | 159 (80%) | 6.4 | 17 |
| gCetuximab | GN451, F1-1 | 129 (65%) | 5.2 | 16 |
| gCetuximab | GN451, F1-2 | 126 (63%) | 5.0 | 22 |
| Erbitux | Commercial | 131 (66%) | 5.3 | 6 |

<sup>a</sup> Relative galactosylation is the sum of the percentage of each carbohydrate structure in the sample multiplied by the number of gal residues in that structure. Since each structure can have one or two gal residues this value can be a maximum of 200. Dividing this number by two gives the percent galactosylation.
<sup>b</sup> Gal/mole is calculated by multiplying the percent galactosylation by eight, the maximum possible number of Gal residues per molecule.

### Binding to EGFR on HTB-30 cells

To be useful as a therapeutic, it was necessary to show that gCetuximab would bind to the intended target EGFR with similar efficiency as Erbitux. A study was performed by comparing the binding of Erbitux and goat-produced antibody to HTB-30 cells, a human breast cancer cell line known to express both EGFR and Her-2 antigens (28). For this assay, serial dilutions of fluorescently labelled antibody were applied to cells and binding to EGFR determined by measuring cell-bound fluorescence. Plotting of the mean fluorescent intensity (MFI) values confirmed that gCetuximab and Erbitux have very similar binding efficiencies (Fig. 7A).

**Fig. 7.**
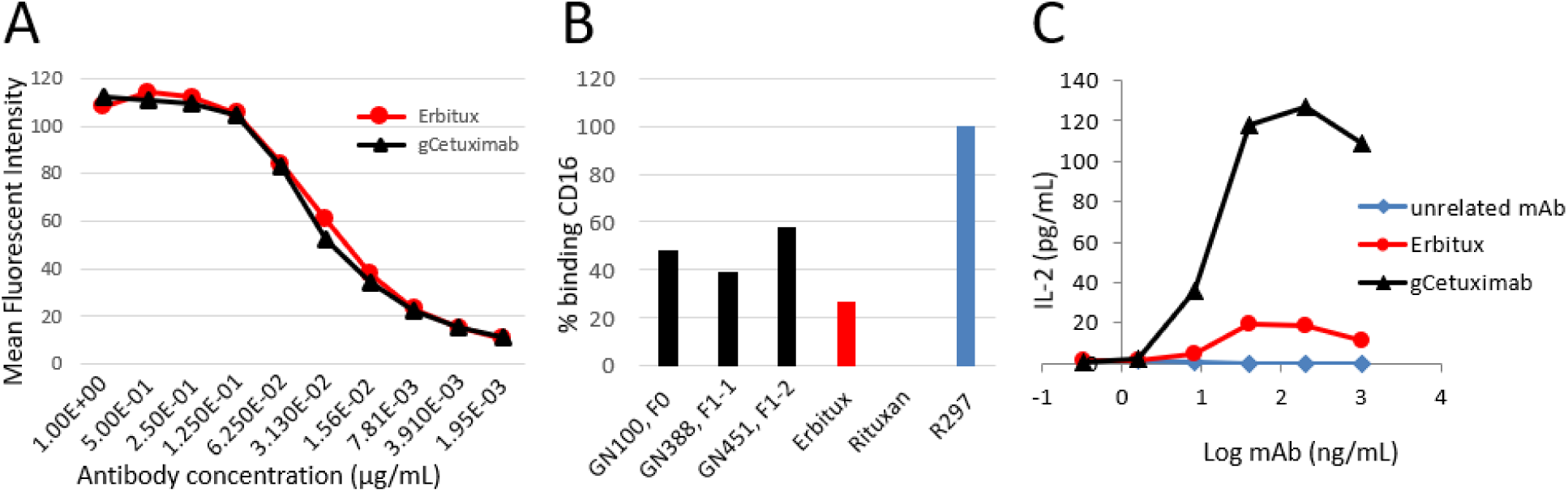
Comparative binding efficiencies of gCetuximab and Erbitux to EGFR and CD16 and their potential for enhanced ADCC. A) Binding efficiency of gCetuximab (black) and Erbitux (red) to EGFR was assessed using flow cytometry by binding serial dilutions of antibody (as indicated) to EGFR-expressing HTB-30 cells. Shown are MFI measurements plotted against antibody concentration to determine binding efficiency. B) CD16 binding on Jurkat-CD16-cells for 3 different batches of gCetuximab produced by F0 and F1 goats from different transgenic lines (black, as indicated), compared to commercial Erbitux (red). The MFI observed are expressed in percent of CD16 binding, with 100% being the value obtained with the R297 anti-D mAb (EMABling, LFB; blue) and 0% the value in the presence of Rituxan. C) Anti-EGFR antibody-bound target cell mediated IL-2 secretion from activated Jurkat-CD16 cells. Shown are the results of two independent measurements of the IL-2 levels in cell culture supernatants for co-cultures of Hep-2 cells incubated with the indicated concentrations of antibodies in the presence of Jurkat-CD16 cells. Unrelated antibody (blue): mAb that does not recognize EGFR; Erbitux (red): commercial cetuximab; gCetuximab (black): purified from F1-2, line GN451.

### Enhanced CD16 binding and improved ADCC

Cetuximab is known to recruit cytotoxic effector immune cells against tumor cells expressing EGFR, resulting in ADCC. To examine whether gCetuximab might exhibit increased ADCC compared to Erbitux we first determined each protein’s ability to bind to CD16, a crucial receptor involved in ADCC. gCetuximab purified from the three different milk samples (GN100, F1; GN388, F1-1; GN451, F1-2) were tested for CD16 binding in competition with a fluorescently labelled antibody. In this assay high fluorescence from high levels of bound competitor indicates low CD16 binding affinity of the tested antibody and conversely, low fluorescence levels a high CD16 binding affinity. MFI values for the three goat-produced antibodies (25.1, 28.1 and 21.8) were lower than for Erbitux (32.4). Binding to CD16 was calibrated against known antibodies with low (Rituxan) and high (R297) CD16 binding affinity with their MFI 41.5 and 7.3 arbitrarily set at 0% and 100% CD16 binding, respectively. All three batches of gCetuximab showed enhanced binding to CD16 (39.3 % – 57.7 %) compared to commercial Erbitux determined at 26.7 % (Figure 7B).

Next, an ADCC bioassay was utilized based on the activation of Jurkat-CD16 cells by antibody-bound target cells. For the assay, Hep-2 cells were utilized as suitable target cells after confirming expression of EGFR on their cell surface (Figure S8).

In the presence of Jurkat-CD16 cells, Hep-2 target cells mediated interleukin-2 (IL-2) secretion from Jurkat-CD16 cells dependent on their anti-EGFR antibody loading (Figure 7C). IL-2 secretion with gCetuximab, purified from F1-2 of line GN451, had a maximal efficacy (Emax) of 127 (±7.8) pg IL-2/mL and a half maximal effective concentration (EC50) of 10 ng/mL for gCetuximab. The equivalent IL-2 secretion mediated by commercial Erbitux was much lower with an Emax value of only 20 (±2.1) pg IL-2/mL. Due to the low secretion levels, we were unable to determine the EC50 value for Erbitux.

## DISCUSSION

Dairy animals represent an attractive alternative production platform for therapeutic proteins compared to prevailing mammalian culture systems. However, only two therapeutic proteins isolated from the milk of transgenic animals have so far entered the marketplace (7-9). They were recently joined by a third product, human coagulation factor VII, that has now gained approval by the FDA (11).

There are a growing number of efficacious mAbs, and the need for large amounts of such therapeutic mAbs for the treatment of serious human diseases. This makes mAbs ideal candidates for production in transgenic dairy animals due to their ability for flexible, economic production of large amounts of mAbs in milk (6, 12, 29) Described here are the first steps to developing an alternative to traditional production of commercial mAbs such as Erbitux. Evidence is presented that stable lines of founder transgenic animals can be obtained that produce high levels of recombinant mAb in their milk. These lines of animals carry transgenes that are stably integrated and were shown to be free of any unwanted mutations.

Next generation sequencing enabled the identification of the chromosomal insertion sites which allowed the confirmation of stable transmission of the transgene copies into the next generation. Hence, conventional breeding approaches can be readily applied for the large scale expansion of the mAb-producing goat lines for commercial gCetuximab production. However, we observed that not all transgenic lines were suitable for large-scale production. Goat lines with integrations of many transgene copies failed to properly lactate, most likely due to high transgene expression levels that might have interfered with the efficient secretion of the mAb and other components naturally found in milk. By contrast, low copy lines with between one and four copies of the LC- and HC-transgenes lactated normally. Overall, no other phenotype was observed, and all goats were healthy.

The high level of production of 5-15g/L compares with the expression level of an anti-CD20 mAb in cattle (12). With the expected goat milk yield of 800L/year, these animals can easily produce multiple kilograms of gCetuximab annually. The mAb is readily accessible in the milk and, as shown in our study, can be efficiently purified, fulfilling an important prerequisite for commercial production.

The monoclonal produced, gCetuximab, has the required equivalent capabilities as the innovator product, Erbitux. Critically, it binds to the EGFR found on tumor cells with comparable affinity to the commercial product and can be efficiently purified, necessary for commercial production.

Significantly, the goat-produced mAb, gCetuximab, has two distinct advantages over the current Erbitux product. First of all, the goat mammary gland does not modify gCetuximab with the highly immunogenic α-Gal epitope. By contrast, α-Gal is present on the current Erbitux product which has been shown to elicit an adverse response in some patients triggered by an immune reaction against α-Gal (14). This would increase the safety profile of treatments with gCetuximab compared to Erbitux.

Secondly, evidence is provided that gCetuximab has increased effectiveness compared to the current Erbitux product. The mechanism of action of Erbitux is believed to be primarily through interference of the binding of EGF to the extracellular domain of the EGFR, blocking dimerization and activation of the receptor (30-32).

ADCC could provide a mechanism to enhance the effectiveness of Erbitux but has not been exploited. This capability of ADCC has recently been recognized by glycoengineering cetuximab for a tighter binding to CD16 associated with increased ADCC (33). The fact that the gCetuximab samples show increased binding to CD16 suggests that they have increased ADCC activity. This was then directly demonstrated with mAbs produced by the goat line GN451, which had the highest CD16 binding affinity. ADCC acitivity of the gCetuximab samples showed a more than six-fold increase compared to the commercial product.

In a separate study, detailed comparative characterisation of gCetuximab vs Erbitux, including *in vitro* and *in vivo* assessments of EGFR inhibition and anti-tumour effects further demonstrated that gCetuximab has the required equivalent capabilities as the innovator product, Erbitux (34).

Taken together, this suggests that gCetuximab not only has similar functionality but could also provide a more effective treatment option compared to Erbitux. Thus, the generation of these lines of animals, capable of high level production, could provide a a cost-effective source of a new and improved version of cetuximab, one without the α-Gal issue as well as increased effectiveness due to enhanced ADCC.

## Abbreviation

ADCC: antibody-dependent cell-dependent cytotoxicity
α-Gal: galactose-α-1,3-galactose
BP: breakpoints
*CEMIP*: cell migration inducing hyaluronan binding protein gene
ddPCR: droplet digital polymerase chain reaction
EC50: half maximal effective concentration
EGFR: epidermal growth factor receptor
Gal: galactose
gCetuximab: goat-produced cetuximab
g*CSN2*: goat β-casein gene
GFF: goat fetal fibroblast
HC: heavy chain
IL-2: interleukin-2
LC: light chain
*LGB*: beta-lactoglobulin gene
mAb: monoclonal antibody
MFI: mean fluorescent intensity
PGK: 3-phosphoglycerate kinase
REF: integrative genome viewer
SCNT: somatic cell nuclear transfer
WT: wild type
*ZNF451*: zinc finger protein 451 gene

## Acknowledgements

The authors kindly thank Stephanie Delaney for ultrasound scanning, Aaron Malthus for milking goats and Ali Cullum and Ruakura farm staff for dedicated animal husbandry; Jan Oliver and Fleur Oback for assistance with SCNT; and David Ashline and Vernon Reinhold of Glycan Connections of Fremont, NH for their contributions in carbohydrate analysis.

This work was funded by the Ministry of Business, Innovation and Employment, AgResearch and LFB-USA.

## Author Contribution

G. Laible, D.N. Wells, W.G. Gavin and H.M. Meade participated in the conception, design, analysis and interpretation of data and obtaining funding; S. Cole, B. Brophy, P. Maclean, D.N. Wells, L.H. Chen, D.P. Pollock, L. C, N. Fournier, C. De Romeuf, N.C. Masiello performed research; G. Laible supervised the project and wrote the MS with contributions from D.P. Pollock, H.M. Meade and revisions from D.N. Wells and W.G. Gavin.

## Competing interests

GL, SC, BB, PM, DNW are employees of AgResearch, LHC, DPP, NCM, WGG, HMM and NF, CDR are employees of LFB-USA and LFB Biotechnologies, respectively. All these organisations have a commercial interests or potential commercial interests in the production of gCetuximab. LC has no conflict of interest or financial conflicts to disclose.

## Notes

### Summary of Updates

updated reference 34

