## Supplemental figures and tables for "Transgenic goats producing an improved version of cetuximab in milk" (from another version of this preprint: version 1, 2020-06-07)

### Supplemental Figures S1 – S8

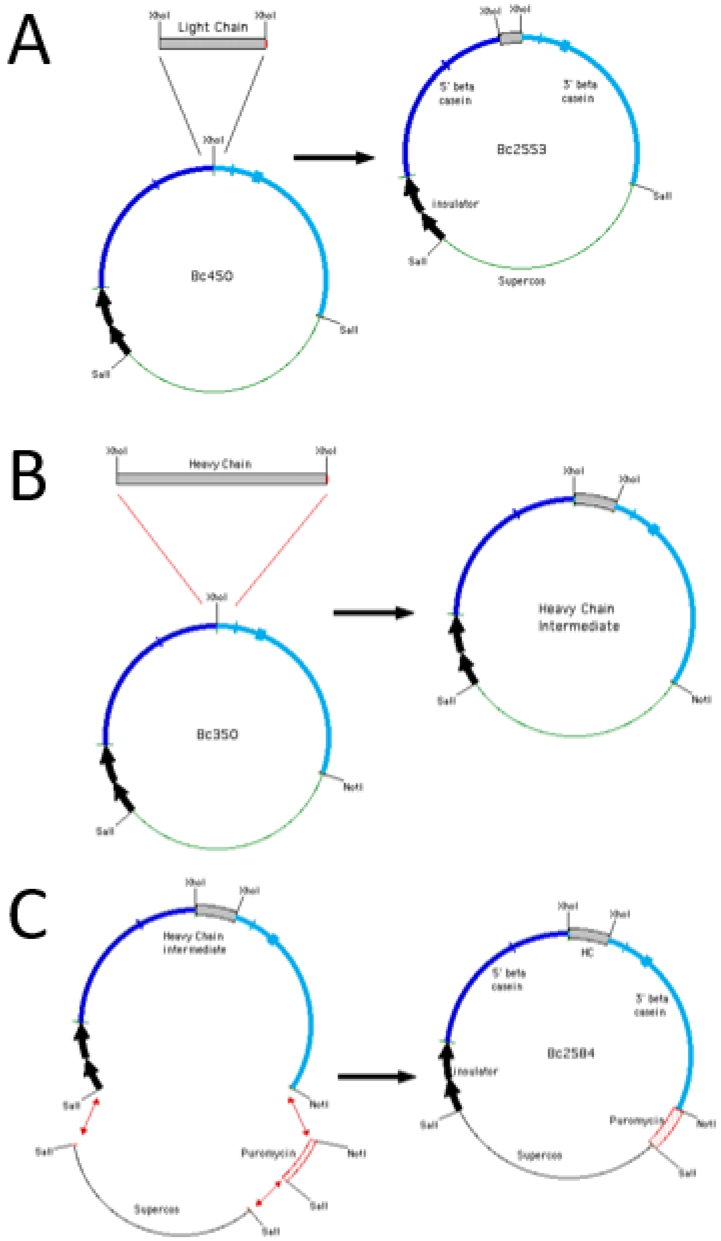


**Fig. S1.** Schematic overview of the construction of the cetuximab expression vectors. A) Light chain expression vector (Bc2553). Tandem black arrows: chicken β-globin insulator sequences; heavy dark and light blue lines: 5’ and 3’ regulatory sequences of g*CSN2*, respectively; light green line: Supercos vector backbone. B) First step in the construction of the heavy chain expression vector generating an intermediate plasmid. C) Second step to generate the heavy chain expression vector. Double headed red arrows indicate correct alignment of compatible restriction sites for successful ligation of three fragments that generated Bc2584.

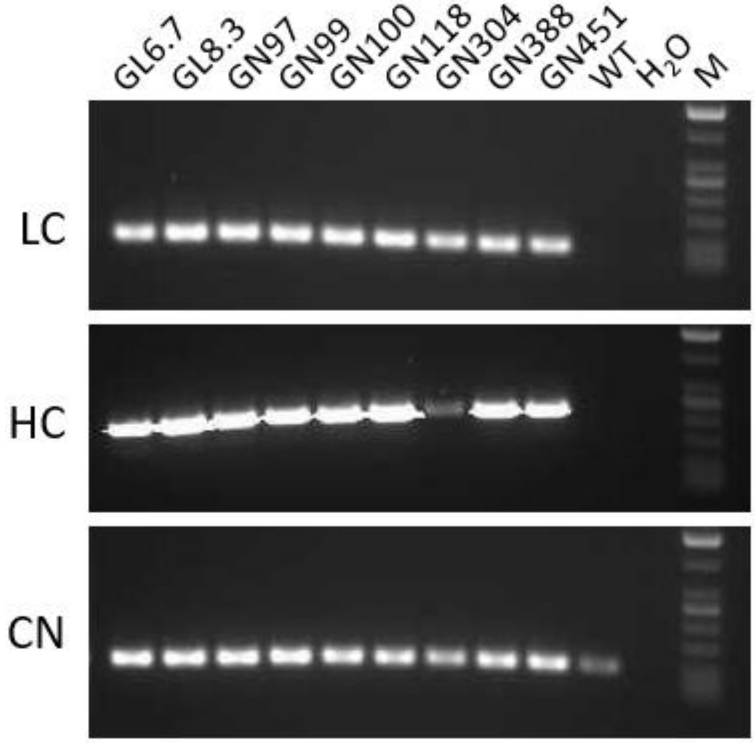


**Fig. S2.** Representative gel photo for the PCR detection of LC- and HC-transgenes in transgenic founder kids for individual lines as indicated. WT: wild type control, $H_2O$: water control, M: size marker, LC: light chain, HC: heavy chain, CN: positive g*CSN2* control.

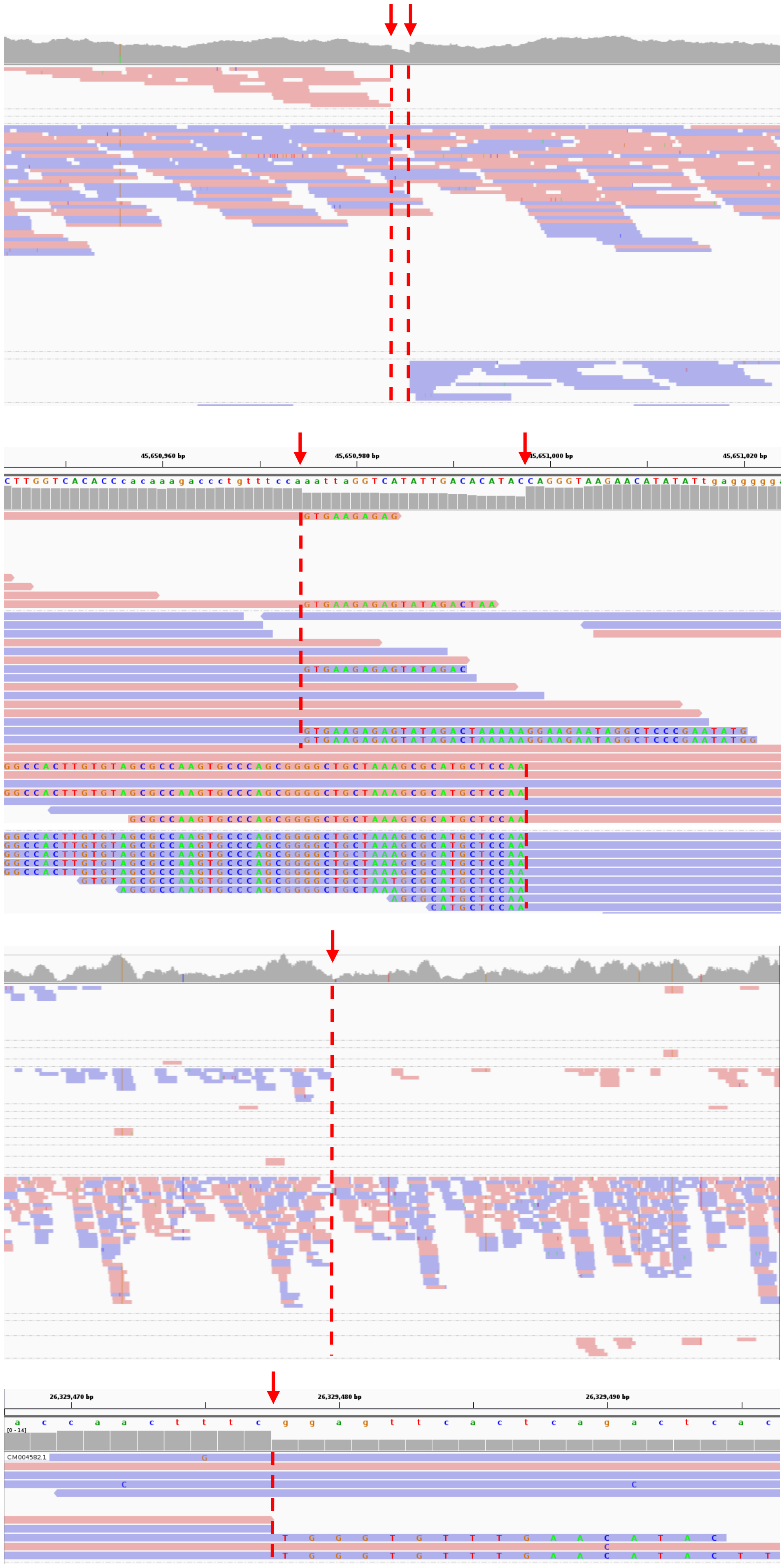

45,650,960 bp
45,650,980 bp
45,651,000 bp
45,651,020 bp
26,329,470 bp
26,329,480 bp
26,329,490 bp
[0 - 14]
CM004582.1

**Fig. S3**. Identification of transgene insertion sites from whole genome sequencing data. A) Screen capture from integrative genome viewer (REF) of the mate pair overlapping the GN388 genomic breakpoints (Chromosome 23 (CM004584.1): 45,650,975 and 45,650,997). The red vertical lines indicate the sites of the breakpoint in the genome where transgenes are inserted. Orange rectangles indicate reads mapping 5' to 3' on Chromosome 23, blue rectangles indicate reads mapping 3' to 5' on Chromosome 23. Coverage of reads mapping to Chromosome 23 is shown in grey at the top. Below are separate groups of reads with mates mapping to different chromosomes or sequences. The mates for the group of orange coloured sequence reads map to CM004567.1 (Chromosome 6) around the CSN2 5' end. The group with orange and blue rectangles depicts reads with both mates mapping to Chromosome 23 and the mates for the blue coloured sequence reads map downstream of the PGK promoter. The high concentration of reads with mates mapping to different chromosomes indicate the general location of an insertion site (red arrows). B) REF screen capture of the paired end reads overlapping the GN388 genomic breakpoints of the transgene insertion site on Chromosome 23 (CM004584.1). The top depicts coverage of mapped sequence reads to the relevant region on Chromosome 23, including the nucleotide sequence of the genomic locus. The red arrows indicate the exact site of the breakpoints on the genome. The nucleotides shown for individual reads indicate mismatches between the reads and reference genome. Nucleotides starting from the left-hand side of the screen capture match the g*CSN2* promoter sequence and the nucleotides starting from the right-hand side of the screen capture match the sequence downstream of the PGK promoter. C) REF screen capture of the mate pair reads overlapping the GN451 genomic breakpoint (Chromosome 21 (CM004582.1): 26,329,477). The red vertical line indicates the 5' end of breakpoint in relation to the genome where transgenes are inserted. Orange rectangles indicate reads mapping 5' to 3' on Chromosome 21, blue rectangles indicate reads mapping 3' to 5' on Chromosome 21. Coverage of reads mapping to Chromosome 21 is shown in grey (top) and followed by separate groups of reads with mates mapping to different chromosomes or sequences. The group of blue rectangles represents mate sequence reads that map to the chicken genome (insulator sequences of the transgenes). The next group of (mainly) blue and orange rectangles indicates mate sequence reads that map to Chromosome 6. The blue coloured reads have their mate mapped around the CSN2 promoter, while the orange coloured have their mate mapped around the CSN2 5' end. The group with intermixed blue and orange rectangles depicts reads with both mates mapping to Chromosome 21. The group of orange coloured sequence reads at the bottom, has mates that map downstream of the PGK promoter. The high concentration of these reads with mates mapping to different chromosomes indicate the general location of an insertion site (red arrow). D) REF screen capture of the paired end reads overlapping the GN451 genomic breakpoint on Chromosome 21 (CM004582.1) where transgenes are inserted. The top depicts coverage of mapped sequence reads to the relevant region on Chromosome 21, including the nucleotide sequence of the genomic locus. The red arrow indicates the exact site of the 5' breakpoint on the genome. The nucleotides shown indicate mismatches between the reads and reference genome. These nucleotides match the 3' arm goat CSN2 sequence of the transgene.

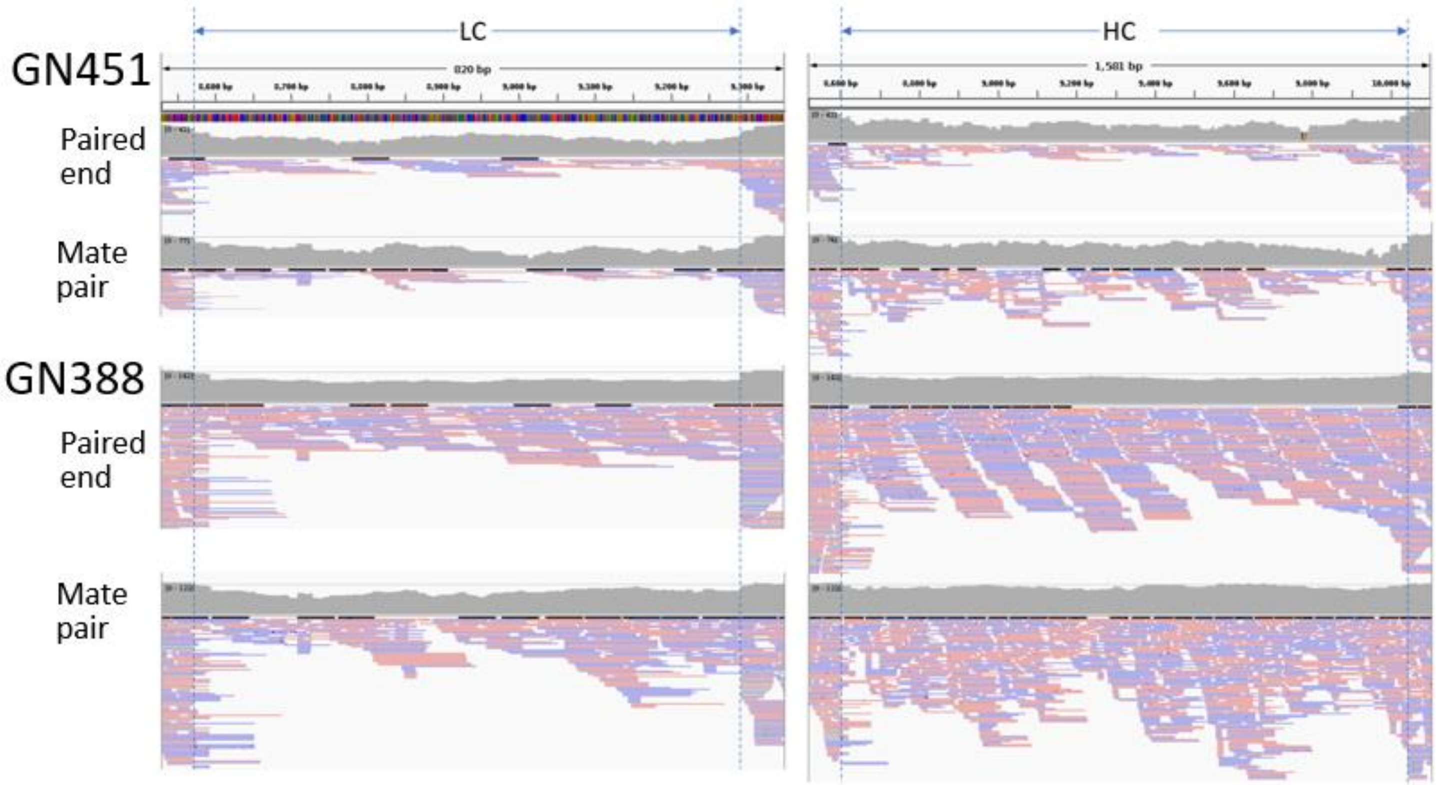


**Fig. S4**. REF screen capture of the coding regions of the LC- and HC-transgenes. Red colored reads are mapped in pairs with read one in forward orientation and read 2 in reverse orientation. Purple colored reads are mapped in pairs with read one in reverse orientation and read 2 in forward orientation. Grey reads indicate that one of the reads in the pair was unable to be mapped. Vertical dashed lines indicate the approximate boundaries of the coding regions of the transgenes. Dashes in the reads indicate mismatches between the read and the transgene sequences. The coverage graph at the top of each panel is on the log scale.

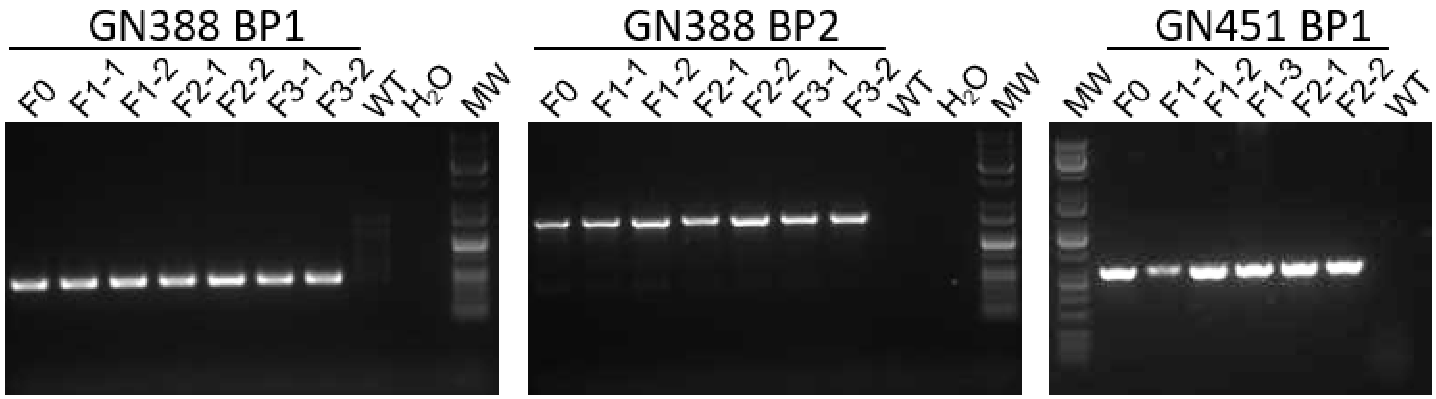


**Fig. S5.** PCR amplification of breakpoints for multiple generations. Shown are fragments amplified by junction PCR across the indicated breakpoints (BP1, BP2) in lines GN388 and GN451. F0: founder; F1-1, F1-2, F1-3: different F1 animals; F2-1, F2-2: different F2 animals; F3-1, F3-2: different F3 animals; WT: wild type; $H_2O$: water control; MW: molecular weight marker.

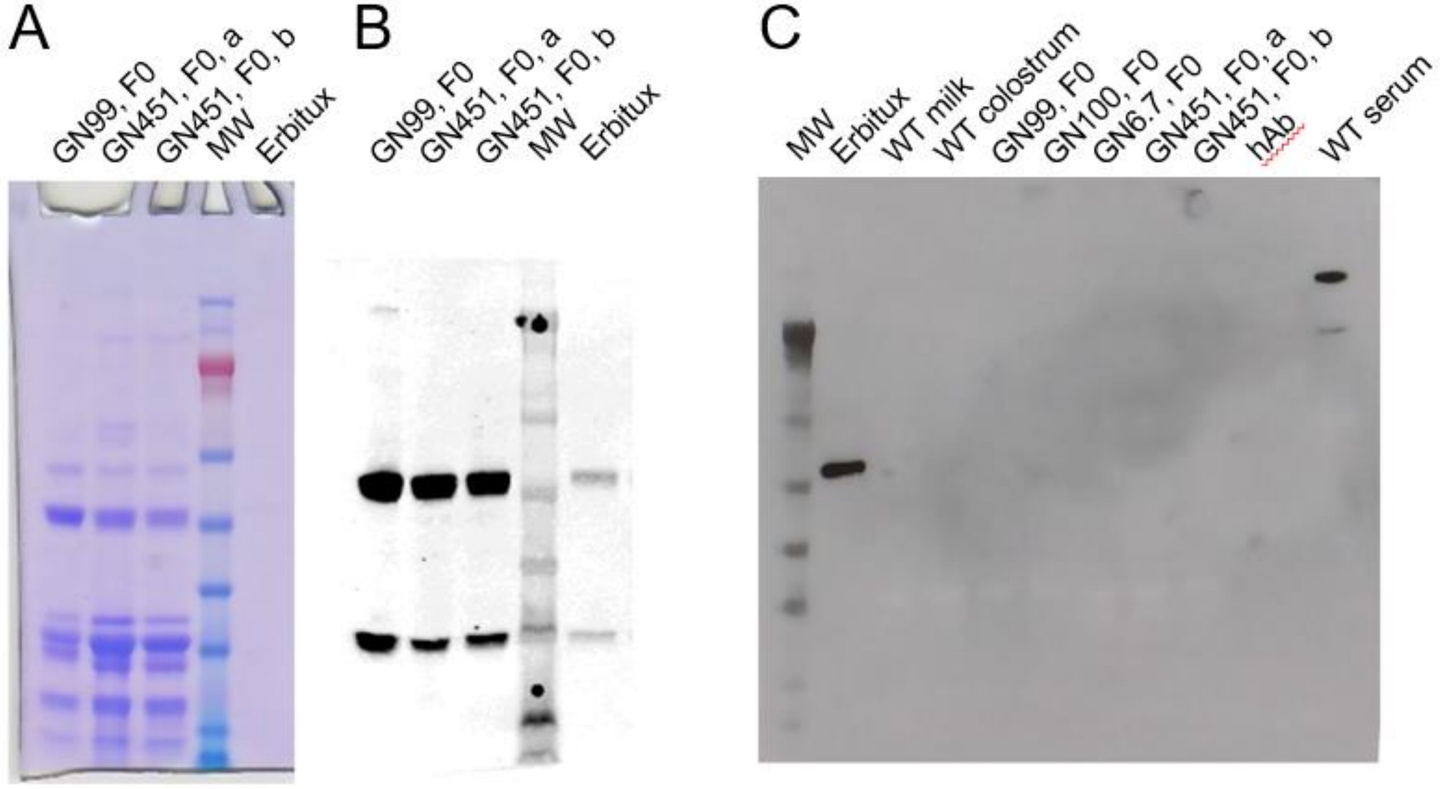


**Fig. S6.** Western blot to confirm lack of α-Gal on gCetuximab. Protein samples were separated under reducing condition by gel electrophoresis and visualized by Coomassie blue staining (A). Western blots were performed to identify human antibodies (B) and assess modification of the proteins by α-Gal (C). Milk, colostrum, and serum samples were all loaded by constant volume of 0.1 µL each, and 300 ng of Erbitux and 1000 ng of an unrelated purified human antibody (hAb) were loaded as controls. MW: molecular weight marker Seeblue +2; Erbitux: commercial cetuximab; WT milk: wild type goat milk; WT colostrum: wild type goat colostrum; GN99, F0: milk from founder goat of line GN99; GN100, F0: milk from founder goat of line GN100; GN6.7, F0: milk from founder goat of line GL6.7; GN451, F0, a and b: two individual milk samples, separated by 12 days, from the founder goat of line GN451; hAb: purified human control antibody; WT serum: wild type goat serum.

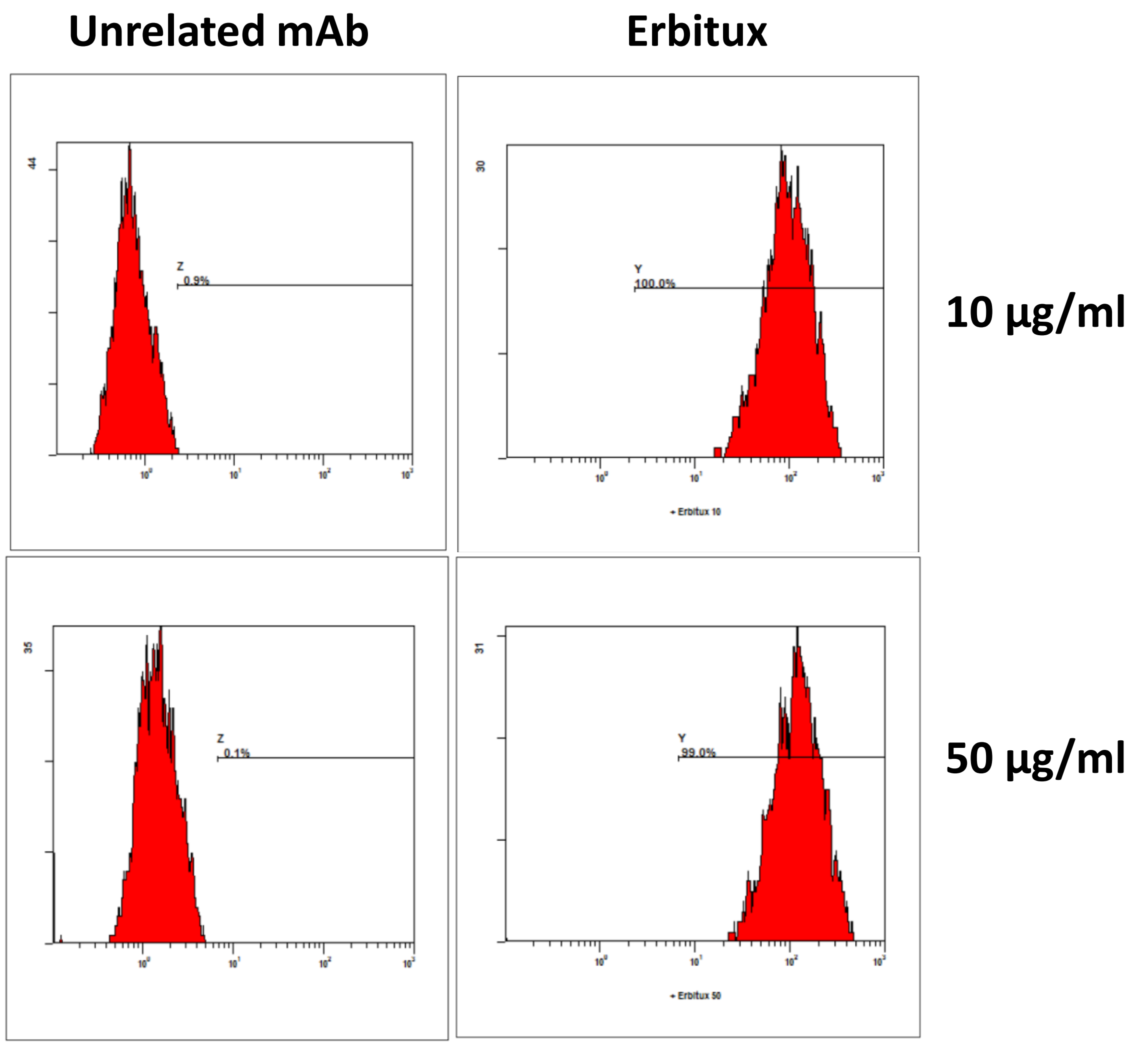


**Fig. S7.** EGFR expression was shown on Hep-2 cells by flow cytometry.

Supplemental Tables S1-S2

**Table S1.** Number of integrated transgene copies in isolated cell clones

| Cell clone | LC copies | HC copies |
|---|---|---|
| GL6.7 | 1 | 2 |
| GL8.3 | high[a] | high[a] |
| GN97 | 8 | 2 |
| GN99 | high[a] | 2 |
| GN100 | high[a] | 2 |
| GN118 | high[a] | high[a] |
| GN304 | 4 | 1 |
| GN306 | high[a] | high[a] |
| GN307 | ND[c] | ND[c] |
| GN362 | high[a] | med high[b] |
| GN363 | high[a] | high[a] |
| GN368 | ND | ND |
| GN369 | 6 | 0 |
| GN381 | 4 | 1 |
| GN383 | high[a] | high[a] |
| GN388 | 2 | 4 |
| GN399 | ND[c] | ND[c] |
| GN400 | high[a] | high[a] |
| GN450 | 1 | 0 |
| GN451 | 1 | 1 |
| GN452 | med high[b] | med high[b] |
| GN453 | high[a] | high[a] |
| GN455 | >8 | 1 |

[a] Copy numbers estimated as >100 were classified as high and not accurately quantified.

[b] Copy numbers estimated to be in the range of 50-100 copies were classified as med-high and not accurately quantified.

[c] Copy numbers could not be determined due to insufficient DNA.

**Table S2.** Transgene copy numbers for A) line GN388 and B) line GN451 as determined by WGS data.

**A**

| **Genomic region** | **Average coverage** | | **Haploid genome coverage** | | **Target genes** | **Estimated copy number** | |
|---|---|---|---|---|---|---|---|
| | Paired end | Mate pair | Paired end | Mate pair | | Paired end | Mate pair |
| **Genome** | 28.1 | 13.7 | 14.0 | 6.9 | Endogenous genes | 2 | 2 |
| **g*CSN2* coding region** | 32.0 | 14.0 | | | g*CSN2* | 2 | 2 |
| **TGs, combined** | | | | | | | |
| 5’ TG flank (g*CSN2*) | 177.2 | 67.9 | | | g*CSN2* + LC + HC | 12 (10)* | 10 (8)* |
| 3’ TG flank (g*CSN2*) | 178.8 | 62.7 | | | g*CSN2* + LC + HC | 13 (11)* | 9 (7)* |
| **TGs, individually** | | | | | | | |
| LC | 20.1 | 15.2 | | | LC | 1 | 2 |
| HC | 43.6 | 35.3 | | | HC | 3 | 5 |
| LC + HC | - | - | | | LC + HC | 4 | 7 |

* brackets indicate total number of transgene copies after the deduction of the two endogenous g*CSN2* copies

**B**

| **Genomic region** | **Average coverage** | | **Haploid genome coverage** | | **Target genes** | **Estimated copy number** | |
|---|---|---|---|---|---|---|---|
| | Paired end | Mate pair | Paired end | Mate pair | | Paired end | Mate pair |
| **Genome** | 14.2 | 13.6 | 7.1 | 6.8 | Endogenous genes | 2 | 2 |
| **g*CSN2* coding region** | 15.3 | 14.6 | | | g*CSN2* | 2 | 2 |
| **TGs, combined** | | | | | | | |
| 5’ TG flank (g*CSN2*) | 27.7 | 23.2 | | | g*CSN2* + LC + HC | 4 (2)* | 3 (1)* |

| | | | | | |
|---|---|---|---|---|---|
| 3’ TG flank (g*CSN2*) | 31.6 | 22.5 | g*CSN2* + LC + HC | 4 (2)* | 3 (1)* |
| **TGs, individually** | | | | | |
| LC | 5.2 | 6.5 | LC | 1 | 1 |
| HC | 4.2 | 9.3 | HC | 1 | 1 |
| LC + HC | - | - | LC + HC | 2 | 2 |

* brackets indicate total number of transgene copies after the deduction of the two endogenous g*CSN2* copies
